# PoMo: An Allele Frequency-based Approach for Species Tree Estimation

**DOI:** 10.1101/016360

**Authors:** Nicola De Maio, Dominik Schrempf, Carolin Kosiol

## Abstract

Incomplete lineage sorting can cause incongruencies of the overall species-level phylogenetic tree with the phylogenetic trees for individual genes or genomic segments. If these incongruencies are not accounted for, it is possible to incur several biases in species tree estimation. Here, we present a simple maximum likelihood approach that accounts for ancestral variation and incomplete lineage sorting. We use a POlymorphisms-aware phylogenetic MOdel (PoMo) that we have recently shown to efficiently estimate mutation rates and fixation biases from within and between-species variation data. We extend this model to perform efficient estimation of species trees. We test the performance of PoMo in several different scenarios of incomplete lineage sorting using simulations and compare it with existing methods both in accuracy and computational speed. In contrast to other approaches, our model does not use coalescent theory but is allele-frequency based. We show that PoMo is well suited for genome-wide species tree estimation and that on such data it is more accurate than previous approaches.

---

Understanding the speciation history of taxa is fundamental to the study of evolution, but is often hampered by incongruency among phylogenetic trees from different genomic regions. Three different biological processes can cause this pattern: horizontal gene transfer, gene duplication and loss, and incomplete lineage sorting (ILS, see Maddison 1997; Knowles 2009). Horizontal gene transfer plays a major role in bacterial evolution, and gene duplication and losses are common throughout the tree of life. The third process, ILS, has received considerable attention from a theoretical point of view, in particular in recent years (see e.g., Maddison and Knowles 2006; Degnan and Rosenberg 2009; Edwards 2009; Liu et al. 2009a). ILS occurs when the coalescent time between two lineages within a branch of the species tree is longer than the branch itself. Simple and computationally efficient approaches such as concatenation of the gene alignments to one overall alignment to infer the species tree (see Gadagkar et al. 2005) or “democratic vote” between gene trees (Pamilo and Nei 1988), proved to be unsatisfactory in accounting for ILS. In fact, they tend to infer the wrong topology for species trees with parameters within the so-called “anomaly zone” as more data are considered (see Degnan and Rosenberg 2006; Kubatko and Degnan 2007).

Several alternative likelihood-based methods have been proposed that explicitly model ILS. These can be divided into two classes: population tree and species tree methods. Although a population tree and a species tree have the same structure, population tree and species tree methods focus on different time scales. In fact, population tree methods model recent population splits by assuming that any polymorphism observed within or between taxa already existed at the phylogenetic root (or equivalently, that there are no new mutations along the phylogeny). Some examples of population tree methods are *δ*a*δ*i (Gutenkunst et al. 2009), and the methods of RoyChoudhury et al. (2008), and Sirén et al. (2011). A partial generalization is SNAPP (Bryant et al. 2012), which allows a single new mutation per site and back-mutations, but still assumes at most two alleles per site. These population tree methods are therefore generally not suited for long phylogenetic distances where two or more mutations are likely to occur at the same site in different points of the phylogeny. Another characteristic of population tree methods is that all sites are modeled as unlinked, ignoring haplotype structures. This can be problematic when few linked loci are considered, but it generally does not lead to biases if many recombining loci are considered, even if linkage within each locus is strong (Wiuf 2006; RoyChoudhury 2011).

Species tree methods, on the other hand, do not rely on the infinite sites assumption, and can generally deal with sites presenting any number of alleles, and arbitrarily long branches. They assume that sites from the same locus (or “gene”) are perfectly linked, while sites from different loci are perfectly unlinked, so that the phylogenetic history of each locus is modeled by exactly one tree (the gene tree) and, conditioned on the species tree and mutational parameters, gene trees are independent of each other. In particular for large samples and high effective recombination rates, these assumptions could also lead to biases in phylogenetic estimation due to intra-locus recombination. Nevertheless, a recent simulation study suggested that these biases are very mild (Lanier and Knowles 2012). Bayesian phylogenetic software for ILS include BEST (Liu et al. 2008), *BEAST (Heled and Drummond 2010), ST-ABC (Fan and Kubatko 2011), and recent versions of MrBayes incorporating the BEST model (Ronquist et al. 2012). Maximum likelihood (ML) methods include STEM (Kubatko et al. 2009) and STELLS (Wu 2012). Other approaches aim to reduce the computational burden of a full-likelihood analysis but still remain in the parametric framework, for example the approximate ML method MP-EST (Liu et al. 2010), but see also GLASS (Mossel and Roch 2010), STAR and STEAC (Liu et al. 2009b), and the approach of Carstens and Knowles (2007). Many nonparametric phylogenetic methods have also been proposed, which are generally faster and do not use sequence data directly, but summary statistics from the gene trees estimated by the user (reviewed in Liu et al. 2009a). Several other phylogenetic software packages solve gene tree incongruencies, but are not tailored in particular for ILS (see e.g., Larget et al. 2010; Boussau et al. 2013), with some exceptions such as DLCoal (Rasmussen and Kellis 2012) which accounts for both ILS and gene duplication and loss.

Recently, we proposed a POlymorphisms-aware phylogenetic MOdel (PoMo) to estimate population genetic parameters (mutation rates and fixation biases) using the genetic variation within and between species (De Maio et al. 2013). In particular, we applied it to exome-wide alignments of 4 great ape species. We included 10 sequences from each species, for a total of 40 sequences and analyzed more than 2 million sites in a single ML estimation run. A brief description of the model will be given below. Here, we extend PoMo to accurately estimate species trees and to overcome discordances in gene trees caused by ILS. PoMo is a species tree method, allowing any number of mutations at each site and branch. Nevertheless, it shares several of its features with many population tree approaches, e.g., all sites are assumed to be unlinked and substitutions are modeled via accumulation of allele frequency changes. With PoMo we aim to overcome the difficulties and caveats associated with parametrizing and estimating gene trees (Knowles et al. 2012). We show that in contrast to other parametric phylogenetic methods, PoMo allows the fast and accurate analysis of large datasets, even genome-wide data. Additionally, our method accounts for within-locus recombination and can easily be used to model fixation biases (such as selection or biased gene conversion) and variation of rates among sites and branches (De Maio et al. 2013).

## METHODS

### Species Tree Inference with PoMo

PoMo is a phylogenetic model of sequence evolution. As in standard models, a single phylogenetic tree (the species tree) represents the speciation history of the considered taxa and all sites are modeled as unlinked. Evolution of a genomic site is modeled as a continuous-time Markov chain along the phylogeny. Yet, in contrast to classical phylogenetic models, PoMo can account for multiple samples from the same taxon. Rather than representing the state of a species with a single nucleotide, PoMo allows species to be polymorphic, with two alleles coexisting at a certain frequency. This is achieved by using a larger state space in the Markov chain. In fact, we use 4 states representing each one of the 4 nucleotides being fixed in the population, and also other states representing polymorphisms. We model evolution of populations in the species tree with a Moran model with population size *N* (called “virtual population” in the following). Hereby we will always use a virtual population size of *N* = 10. The 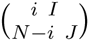 with 1 *≤ i ≤ N −* 1 represents the polymorphic state associated with alleles {*I, J*} and with frequency *i/N* of allele *I*. In a Moran model, the probability of a population changing in one generation 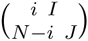 to 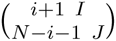 is:

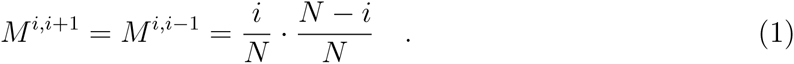

We use this model of genetic drift together with the HKY model of mutation to define an instantaneous rate matrix (see online Appendix 1). This matrix is used in the same way as a standard DNA substitution model. In particular, we use the Felsenstein pruning algorithm (Felsenstein 1981) to calculate likelihoods, summing over all possible histories of mutations and frequency changes. In this work, we do not model fixation biases (e.g., selection) or variation in mutation rates. However, these features have already been introduced in De Maio et al. (2013).

Although a single tree (the species tree) is assumed for all loci and sites, PoMo naturally accounts for ILS, because ancestral species are allowed to be in polymorphic states. For example, let us consider the particular evolutionary history depicted in Figure 1a. Here, an ancestral polymorphism is still present in the species from which sequences B1 and B2 are sampled (none of the two alleles has reached fixation). In this scenario, B1 is more closely related to A1 and A2 than to B2 at the considered site, despite B1 and B2 being in the same species.

**Figure 1:**
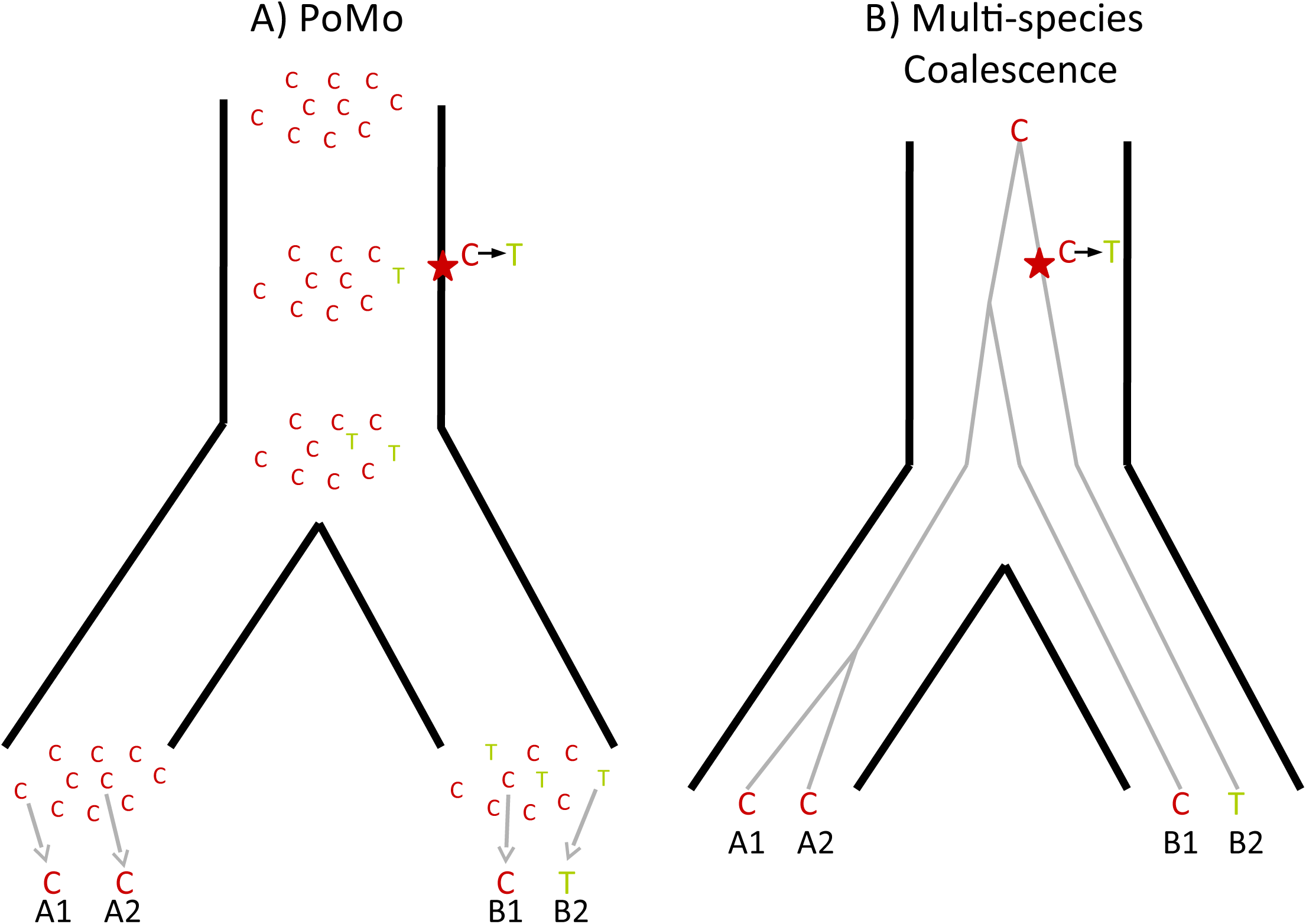
Comparison of PoMo with the multispecies coalescent. Example of a phylogeny with two species, each with two sampled sequences per population (respectively A1-A2, and B1-B2). A single alignment site is considered for simplicity, and the observed nucleotides are as depicted: C, C, C, T. a) In PoMo, observed nucleotides are modeled as sampled from 10 virtual individuals (grey arrows at the bottom). Mutations (stars in the figure) can introduce new alleles. Allele counts can change along branches due to drift, and allele counts can change along branches due to drift, and be lost or fixed. The state history shown is only one of the many possible for the observed data. b) In the multispecies coalescent the species tree (black thick lines) as well as gene trees (grey thin lines) are considered. Usually only the species tree parameters are of interest, and gene trees are nuisance parameters. One of the many possible unobserved gene trees is depicted as an example.

The same situation is also modeled by the multispecies coalescent framework (Rannala and Yang 2003), as can be seen in Figure 1b. In this context, a local phylogeny (the gene tree) models the local relatedness of all samples, and so, here B1 is again less related to B2 than to A1 and A2 at the considered site. In the multispecies coalescent, gene trees are embedded within a species tree, which means that in gene trees a coalescent event between lineages from different species can only happen before (higher up in the tree) the split of the considered species in the species tree. In many applications of the multispecies coalescent, gene trees are modeled as constant within loci and unlinked between loci. In PoMo, on the other hand, all sites, even those within the same gene, are modeled as unlinked.

PoMo has a small number of parameters, and, importantly, the number of parameters does not depend on the number of genes considered or on the number of samples per species. As in most parametric species tree methods we parameterise the species trees (branch lengths and topology). Then, we have parameters that describe the mutational process: in the case of the HKY85 adopted here, these are the mutation rate *μ* and the transition-transversion rate ratio *κ*. Finally one parameter, *θ*, represents the proportion of polymorphic sites, similarly to STEM; yet, differently from STEM, we estimate *θ* from data, and only ask the user to provide an input value for it in case there is a single sample per species, and therefore no information on within-species variation. Differently from *BEAST and BEST, we do not parameterize gene trees, and differently from STEM and MP-EST, we do not ask the user to estimate them, but we implicitly marginalize over all possible histories at each site, and do not make use of haplotype information.

#### Modeling the sampling process in PoMo. —

We introduce a slight modification to PoMo to include the sampling process into the model. In fact, observed nucleotide frequencies do not necessarily match the real population frequencies. If this is not accounted for, sampling variance could be interpreted as drift by PoMo, resulting in biases in branch length estimation. Furthermore, this modification allows the use of any number of samples (even one) for each of the species considered.

Let us assume that at a leaf of the phylogenetic tree allele *I* is observed *m*_*i*_ times, and allele *J* is observed *m*_*j*_ = *M – m*_*i*_ times (*M* being the sample size). We also assume that at the considered leaf the virtual population has count *n*_*i*_ for allele *I* and count *n*_*j*_ = *N – n*_*i*_ for allele *J* (which cannot be observed directly, but we fix it for now). This corresponds to the state represented as 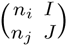 in our Markov chain (see Eq. 1 and online Appendix 1). Each sampled allele is necessarily present in the virtual population (*m*_*i*_ *>* 0 implies *n*_*i*_ *>* 0), that is, we do not model sequencing errors. On the other hand, an allele in the virtual population can be absent from the sample due to chance (*n*_*i*_ *>* 0 but *m*_*i*_ = 0). We use the binomial distribution to model the probability of sampling with replacement *m*_*i*_ times allele *I* and *m*_*j*_ times allele *J* from the virtual population with *n*_*i*_ times allele *I* and *n*_*j*_ times allele *J*:

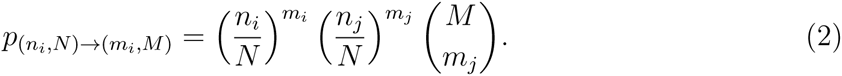

It is straightforward to include this sampling step in the likelihood calculations that are performed with the Felsenstein pruning algorithm (Felsenstein 1981), such that at each leaf we sum over all possible virtual population allele counts.

#### Topology search. —

To efficiently explore the topology space, we included a nearest neighbor interchange (NNI, see Felsenstein 2004) branch swap search in PoMo. Given an unrooted tree ((*T*_1_, *T*_2_), (*T*_3_, *T*_4_)), where *T*_*i*_ for *i* = 1, 2, 3, 4 is a subtree, an NNI move swaps the subtrees to estimate the likelihoods of the alternative topologies ((*T*_1_, *T*_3_), (*T*_2_, *T*_4_)) and ((*T*_1_, *T*_4_), (*T*_2_, *T*_3_)). If any of the two alternative topologies results in a likelihood improvement, it becomes the new base tree for the next NNI step. For every internal branch of the base tree an NNI swap is attempted. The iterations stops if no likelihood improvement is obtained.

## Simulations

Our simulations fit the assumptions of multispecies coalescent methods such as STEM (Kubatko et al. 2009), BEST (Liu et al. 2008), and *BEAST (Heled and Drummond 2010), but not those of PoMo or SNAPP (Bryant et al. 2012). We simulated gene sequences under the standard coalescent model (free recombination between genes, no recombination within genes), each gene being 1kb long. We wanted to address possible differences in performance among methods due to total tree height, tree shape, and sampling strategy (see e.g., McCormack et al. 2009; Huang et al. 2010; Leaché and Rannala 2011; Knowles et al. 2012). We simulated 12 different scenarios according to the species trees depicted in Figure 2. All trees have 4 or 8 species and at least one short internal branch causing ILS. In the “trichotomy” scenario (Fig. 2i) an internal branch has zero length, such that three species are equally related to each other. In the “classical ILS” scenario (Fig. 2ii) an internal branch has short length (*N*_*e*_/10, where *N*_*e*_ is the effective population size), therefore ILS is expected to be predominant. In the “anomalous” scenario (Fig. 2iv) both internal branches are very short, such that the species tree falls inside the “anomaly zone” described by Degnan and Rosenberg (2006). Lastly, for the “recent radiation” scenario (Fig. 2v) all terminal branches, except the outgroup branch, are very short, and different species are expected to share a large proportion of polymorphisms. The last two scenarios include 8 species, and either a balanced topology (“balanced”, Fig. 2iii), or an unbalanced one (“unbalanced”, Fig. 2vi). Each of these 6 scenarios is simulated with total tree height 1*N*_*e*_ and 10*N*_*e*_ generations leading to a total of 12 simulation settings.

**Figure 2:**
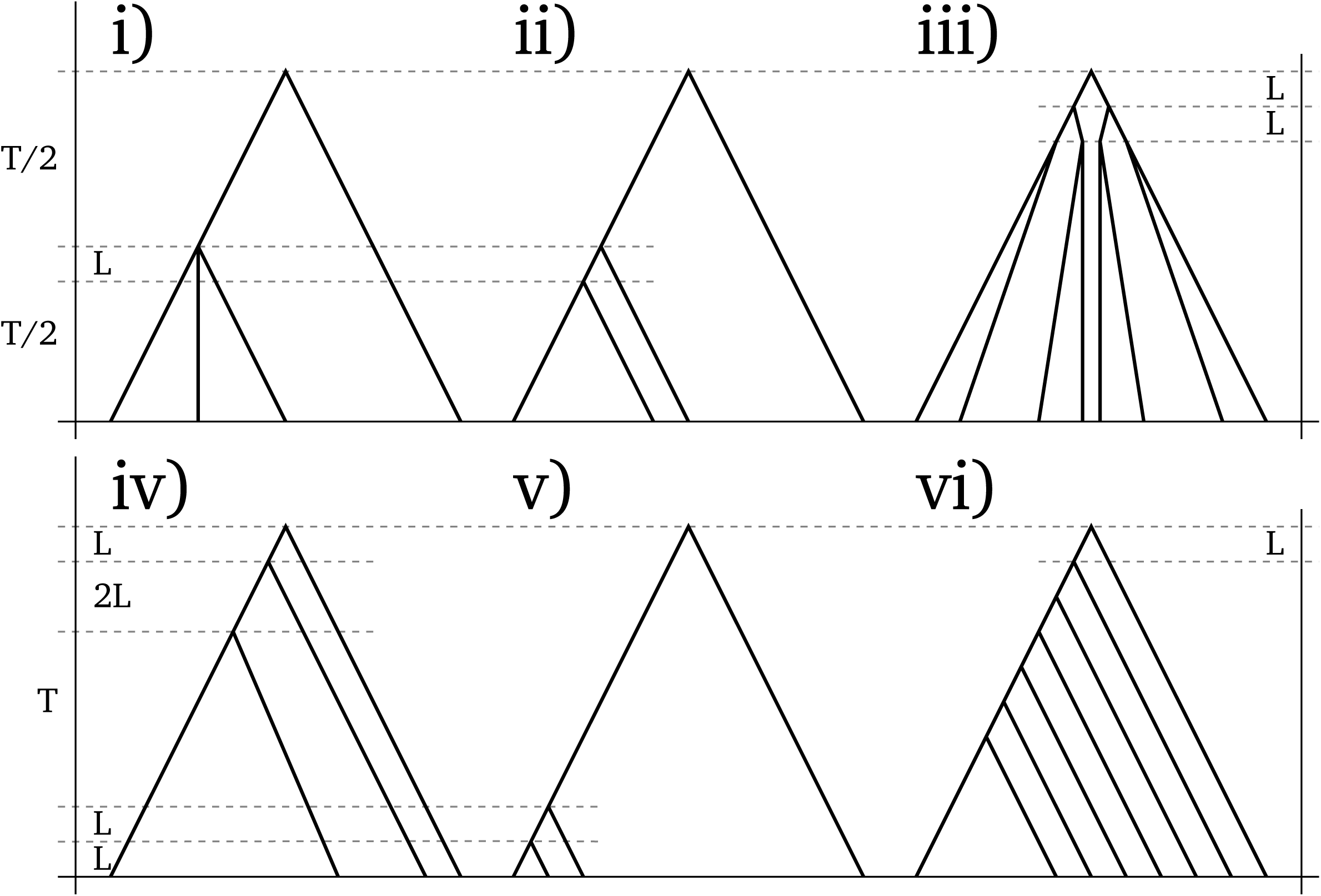
Species Trees used in Simulations. We chose trees that are well-known for inference problems caused by incomplete lineage sorting. Each of the 6 trees shown is used in two scenarios: total tree height (*T*) is either set to 1*N*_*e*_ or to 10*N*_*e*_ generations, where *N*_*e*_ is the effective population size. *L* represents a short branch length of *N*_*e*_/10 generations. Values not shown are determined by the strict molecular clock assumption. The scenario names are i) trichotomy, ii) classical ILS, iii) balanced, iv) anomalous, v) recent radiation and vi) unbalanced.

For each scenario and each replicate we simulate between 3 and 1000 genes. Gene trees from each species tree were sampled according to the multispecies coalescent using MSMS (Ewing and Hermisson 2010). We used three different sampling strategies, extracting either 10, 3, or 1 samples per species. For each gene tree thus simulated, the 1 kbp sequence alignment was generated with Seq-Gen (Rambaut and Grass 1997) according to an HKY model with *к* = 3 and nucleotide frequencies *π*_*A*_ = 0.3, *π*_*C*_ = 0.2, *π*_*G*_ = 0.2 and *π*_*T*_ = 0.3. These settings closely match those in previous simulation studies such as Huang et al. (2010).

This simulation strategy, with trees fixed a priori rather than sampled from a prior distribution, and a single tree as an estimate, might partially favor maximum likelihood methods over Bayesian methods. Also the prior probability of the chosen species trees in the Bayesian methods might play an important role. Yet, it is expected that with increasing amounts of data, the priors will have a smaller and smaller influence on the estimated posterior distribution.

### Comparison of Species Tree Methods

In this work, we compared the performance of PoMo with other methods in estimating species trees. We tested STEM, BEST, and *BEAST, which are all based on the multispecies coalescent equations by Rannala and Yang (2003), and SNAPP, which is instead a population tree method. Furthermore, we also test concatenation, which corresponds to ignoring the effects of ILS and assuming a single phylogeny for the whole dataset. From each simulated dataset and each method we get one estimated species tree, and we assess the accuracy of estimation by comparing the normalized simulated tree and the normalized estimated tree. Normalization is achieved by dividing all branch lengths in the tree by the root height. The normalized trees are then compared using the Branch Score Distance (BSD, see Kuhner and Felsenstein 1994), calculated with TREEDIST from PHYLIP (Felsenstein 2014). BSD uses both topology and branch lengths to assess estimation accuracy, thus providing a broader picture than methods that use only the topology (see also Heled and Drummond 2010). Furthermore, in scenarios where an internal branch is so short as to almost correspond to a trifurcation, for example, (*A* : 1 + *δ,* (*B* : 1, *C* : 1) : *δ*) for *δ →* 0, the BSD has the quality to attribute small error to trees that approach the trifurcation, but have the wrong topology, e.g., (*B* : 1 + *δ,* (*A* : 1, *C* : 1) : *δ*). Below we give a short description of each method tested.

PoMo was implemented in HyPhy (Pond et al. 2005). We explore the topology space using NNI moves without a molecular clock on unrooted topologies using the maximum likelihood of PoMo as a score measure for each topology. We then use the chosen topology to estimate branch lengths with PoMo and maximum likelihood under a strict molecular clock. In contrast to De Maio et al. (2013), we adopt an HKY mutation model (Hasegawa et al. 1985) without fixation biases and without variation in mutation rates. These changes, and in particular the adoption of a molecular clock, have been introduced to fit PoMo to the assumptions of competing approaches and of our simulations. They can easily be reverted by the user.

STEM (Kubatko et al. 2009) is a ML approach. It estimates the ML species trees from a collection of gene trees provided by the user. We estimated gene trees with Neighbor Joining (NJ, see Saitou and Nei 1987) as implemented in HyPhy, and with the Unweighted Pair Group Method with Arithmetic mean (UPGMA, see Murtagh 1984) as implemented in the R package phangorn (Schliep 2011). The input parameter *θ* in STEM is fixed to 0.01.

The program BEST 2.3 (Liu et al. 2008) implements a Bayesian method and, in contrast to STEM, accounts for uncertainty in gene tree estimation. The Felsenstein pruning algorithm (Felsenstein 1981) is used to calculate the likelihood of gene trees, and the multispecies coalescent equation (Rannala and Yang 2003) for the likelihood of the species tree given the gene trees. We stopped the MCMC iteration in BEST when the software’s topological convergence diagnostic reached values below 5%. We did not alter the method’s default priors. As in all other methods tested here, and as in simulations, we used an HKY mutation model. As a species tree estimate we chose the consensus tree with all compatible groups produced by BEST.

*BEAST (Heled and Drummond 2010) is implemented within BEAST2 (Drummond et al. 2012). *BEAST is also a Bayesian method sampling gene trees and species tree with a MCMC approach. An adequate number of MCMC steps has to be specified prior to analysis, and we used two different values: 10^7^ and 10^8^. These values were among the highest possible given our computational resources, but found to be insufficient to achieve convergence in the scenarios with many samples and genes. Therefore, the accuracy and computational cost that we show must always be considered relative to the number of MCMC steps chosen. We allowed no site or locus variation in mutation rates, and no variation in population size matching the assumptions of our simulations. We also used the default priors. We used TreeAnnotator, included in BEAST, to summarize the output of *BEAST into a consensus tree with mean node heights.

SNAPP (Bryant et al. 2012) is also implemented in BEAST2, and is based on an MCMC algorithm that samples species trees from a posterior distribution. Yet, differently from *BEAST and BEST, it does not parameterize gene trees and assumes all sites to be unlinked. Again, we did not alter the default priors of the method. We ran SNAPP for 10^4^ MCMC steps, leading to the same issues as discussed above for *BEAST. No site variation or demographic variation were allowed, and we summarized the posterior distribution with a consensus tree with mean node heights.

Lastly, as representative of ML concatenation we used HyPhy with an NNI branch swap topology search and molecular clock. As a Bayesian representative of concatenation we used MrBayes 3.2 (Ronquist et al. 2012). For both methods we used two concatenation strategies. In the first one we sampled only one individual from each species. In the second strategy for each species we used the consensus sequence of all sampled individuals. As a tree estimate for MrBayes 3.2 we chose the consensus tree with all compatible groups. The full list of options for all software packages used is provided in the Online Appendix 3.

### Great Ape Data Set

The great ape family constitutes one of the most important examples for shared ancestral polymorphisms and ILS (Dutheil et al. 2009), with the species phylogeny comprising variation of evolutionary patterns, closely related taxa and short internal branches.

We used PoMo to estimate evolutionary parameters from exome-wide great ape alignments (*H. sapiens, P. troglodytes, Pon. abelii* and *Pon. pygmaeus*; see De Maio et al. 2013). Here, we modify the dataset to include exome-wide sequencing data from a recent study on the genetic diversity and population history of great apes (Prado-Martinez et al. 2013). The authors provide sequence data of 79 wild- and captive-born individuals including all six great ape species divided into 12 populations. The number of individuals per population ranges from 1 (*Gorilla gorilla diehli*) to 23 (*Gorilla gorilla gorilla*).

We extracted 4-fold degenerate sites from exome-wide CCDS alignments downloaded from the UCSC table browser (

~~~
http://genome.ucsc.edu
~~~

) with *hg18* as the human reference genome (exact download preferences can be found in the Online Appendix 4).

The great ape data on the population level was retrieved from 

~~~
ftp://public_primate@biologiaevolutiva.org/VCFs/SNPs/
~~~

 (Prado-Martinez et al. 2013) in Variant Call Format (VCF). In concatenation analyses we randomly extracted one individual out of each population for each independent run. For PoMo, if more than 10 haplotypes from the same population were present, 10 were randomly sub-sampled. PoMo requires *fasta* format files containing the aligned sequences for all samples, but we wrote a python library (*libPoMo*) to extract data from VCF files. PoMo v1.1.0 was used throughout this paper. We provide documentation of the program (

~~~
http://pomo.readthedocs.org/en/v1.1.0/
~~~

) as well as detailed description of the data preparation and conversion scripts for the great ape data set in the supplementary material (Online Appendix 4).

## Results and Discussion

### Computational Efficiency

Among the approaches that we tested, the most computationally demanding proved to be the Bayesian multispecies coalescent methods: BEST and *BEAST (Figs. 3, S1, S2). Achieving convergence with BEST was beyond our computational resources on many scenarios with as few as 10 genes. The running time of these two methods seems in fact to increase steeply with the number of genes. It might seem that the running time of BEST increases faster than the one of *BEAST, but it has to be considered that we halted BEST when we reached a convergence diagnostic threshold, while we ran *BEAST for a pre-specified number of MCMC steps. BEST and *BEAST are similar in many aspects, in particular they both parameterize explicitly gene trees (with branch lengths and topologies parameters) and explore the space of possible gene trees using MCMC. Therefore, it is not surprising that with an increasing number of genes or samples, the number of MCMC steps required to achieve convergence also increases, as the parameter space to be explored becomes larger. So, on larger datasets, the number of steps that we specified in *BEAST becomes insufficient. Nevertheless, the computational demand of *BEAST increases, as the running time of each MCMC step grows. For these reasons, running Bayesian multispecies coalescent methods such as BEST and *BEAST on datasets with large numbers of loci and samples does not seem feasible.

**Figure 3:**
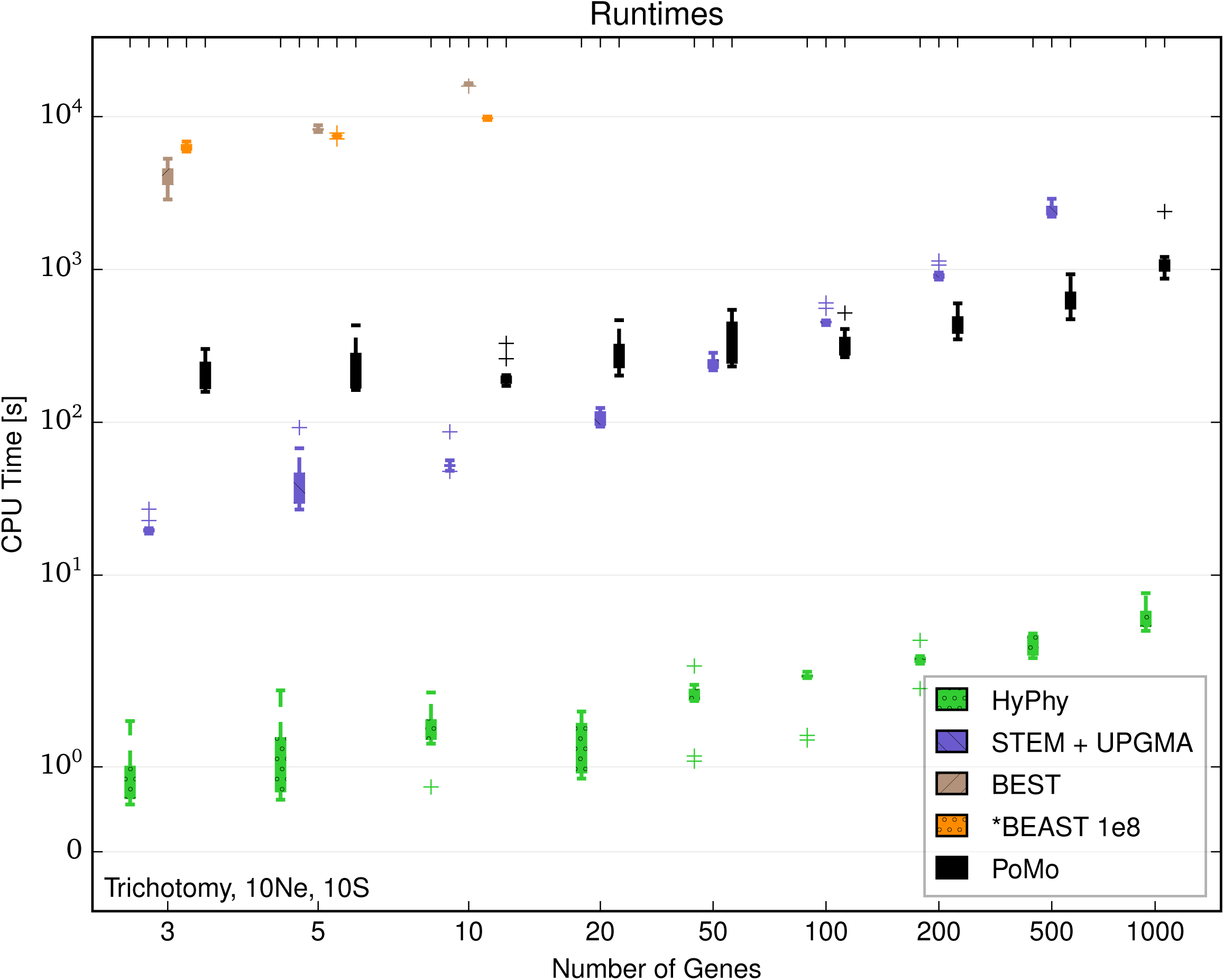
Computational Demands for Different Methods. Running times for estimation with 10 samples per species and tree height 10*N*_*e*_ generations in the trichotomy scenario. The Y axis shows the computational time in seconds, the X axis the number of genes included in the analysis. The colours represent different methods (see legend). Each boxplot includes 10 independent replicates. HyPhy applied to concatenated data is the fastest method. STEM estimates the ML species trees from a collection of gene trees provided by the user. We estimated the gene trees with the Unweighted Pair Group Method with Arithmetic mean (UPGMA) and added the CPU times. For small data sets, PoMo and STEM+UPGMA have comparable computational demands. However, with more genes the CPU time for STEM+UPGMA increases roughly linearly with the number of genes while the time for PoMo remains almost constant. BEST and *BEAST were applied at most to 10 genes. 10^8^ MCMC steps have been used for *BEAST. Our simulations suggest that methods such as *BEAST and BEST are not efficient enough to analyse large datasets.

We ran SNAPP for a very limited number of MCMC steps (10^4^) determined by our computational resources, similarly to *BEAST. Differently from *BEAST, the parameter space of SNAPP does not increase with the number of genes, yet, it still proved computationally demanding, in particular in scenarios with many samples (compare Fig. S5 and S6).

STEM, if considered alone, had extremely short running times, in general only a few seconds (data not shown). Yet, here we consider also the cost of running gene tree estimation, as gene trees are a necessary input for STEM, and they are usually not known. The cost of running gene tree estimation varies greatly depending on the method chosen, but the requirement of ultrametric gene trees (where all leaves have the same distance from the root) restricts the number of possible methods. Here we used two heuristic approaches: Neighbor Joining and UPGMA. These are faster than Bayesian or maximum likelihood methods (UPGMA in particular) and therefore allow the use of STEM even with many genes and samples (Fig. 3). Since the running time of STEM itself is negligible, we see that the computational cost of running STEM and gene tree estimation is approximately a linear function in the number of genes (Figs. S1, S2). Therefore it is feasible to run STEM with exome-wide data, in particular considering that gene tree estimation is easily parallelizable. But, if we would run STEM on an entire eukaryotic genome, for example on millions of loci each of few kbp, then gene tree estimation as performed by us could require months.

In contrast to STEM (which in the following we will always consider including gene tree estimation) PoMo shows a less steep increase in computational requirement with number of genes (Fig. 3). Furthermore, PoMo is almost unaffected by the number of samples included (compare Figs. S1 and S2). A result of this is that while STEM is faster than PoMo on a few genes, with hundreds or thousands of genes PoMo is faster instead, although the particular point at which PoMo becomes faster will depend on the particular gene tree method used and on the number of species and samples considered. Even with 1000 genes, it was always possible to run PoMo in less than an hour with 4 species, or in few hours for 8 species (Figs. S1 and S7). Therefore, PoMo seems well suited for exome-wide species tree estimation, and due to the small increase in running time when adding more data, it is also promising for genome-wide datasets.

Lastly, concatenation methods were generally, but not always, faster than all other approaches (Fig. 3). Concatenation, both Bayesian (with MrBayes) or maximum likelihood (with HyPhy), only allows one sample per species, and has a simplified model (a single tree for all sites) which ignores ILS. It is therefore not surprising that it is faster (Figs. S3 and S4). In particular, the algorithmic steps of PoMo are almost the same as for a maximum likelihood concatenation method, the major differences being the increased dimension of the PoMo rate matrix, and the increased number of internal nodes since PoMo leaf states are not directly observed. Yet, looking at results with 8 species, we see that running times of concatenation on MrBayes are similar to those of PoMo, while running times of concatenation in HyPhy are similar to those of STEM with UPGMA gene trees (Figs. S7 and S8). This suggests, that with more species, faster phylogenetic methods would be required to use concatenation.

We performed all the analyses on the Vienna Scientific Cluster (VSC-2; AMD Opteron 6132 HE processors with 2.2 GHz and 16 logical cores). Every process was assigned to a single core only, so that simulation run times are easily comparable.

### Accuracy in Species Tree Estimation

PoMo shows good accuracy in estimating species tree topology and branch lengths according to the BSD score. While accuracy is variable when only few genes are considered, it rapidly increases with the addition of more data (Figs. 4, 5, S9, S10). Already with 100 genes, errors are below 5%, and get even lower with 1000 genes, although specific values vary with the scenario considered. In fact, errors are usually lower for long species trees (10 *N*_*e*_ generations root height) than for short species trees (1 *N*_*e*_ generations root height). Long trees might be estimated more accurately because they have more phylogenetic signal, or maybe because the contribution of ILS is proportionally less important.

**Figure 4:**
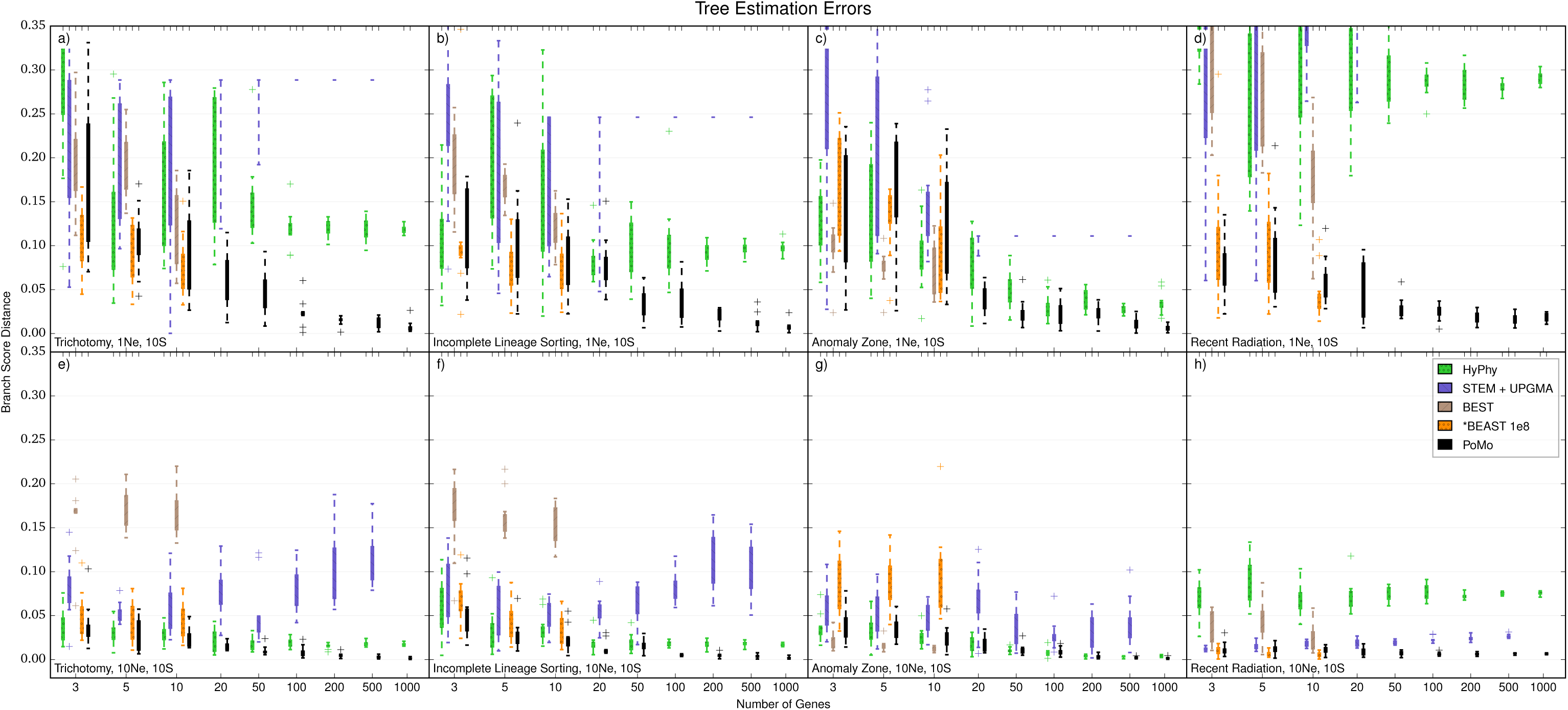
Accuracy of Species Tree Estimation, 4-Species-Trees. We used Branch Score Distance (BSD) to compare the normalized simulated tree and the normalized estimated tree and to measure the error. BSD uses both topology and branch lengths to assess estimation accuracy, providing a broader picture than methods that use only the topology. Higher BSD values indicate larger inference errors. The Y axis is the error in species tree estimation calculated as BSD, the X axis the number of genes included in the analysis. 4 Species and 10 samples per species were included. Each boxplot includes 10 independent replicates. Different colors represent different methods (see legend). a) 1*N*_*e*_ tree height and scenario with trichotomy. b) 1*N*_*e*_ tree height and ILS scenario. c) 1*N*_*e*_ tree height and anomalous species tree. d) 1*N*_*e*_ tree height and recent population radiation. e) 10*N*_*e*_ tree height and trichotomy. f) 10*N*_*e*_ tree height and ILS scenario. g) 10*N*_*e*_ tree height and anomalous species tree. h) 10*N*_*e*_ tree height and recent population radiation. BEST and *BEAST were applied at most to 10 genes. 108 MCMC steps have been used for *BEAST. Note that the alternative methods are often inconsistent, i.e. the error in tree estimation did not decrease as more data was added. PoMo shows accurate parameter estimates that converge towards the true values as more genes are included in the analysis in all scenarios.

**Figure 5:**
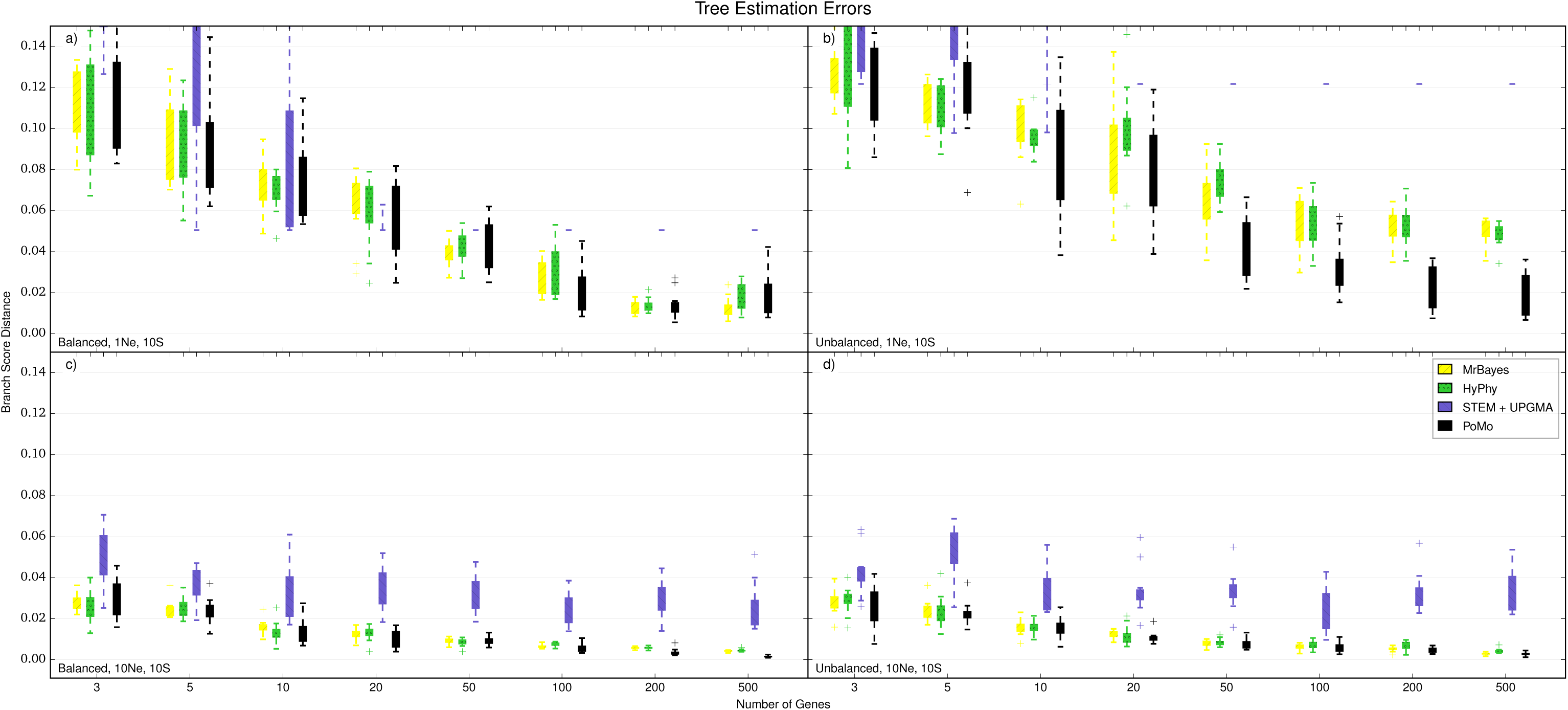
Accuracy of Species Tree Estimation, 8-Species-Trees. For larger trees only PoMo, STEM+UPGMA and concatenation with MrBayes or HyPhy could be used. 8 species and 10 samples per species were included. The Y axis is the error in species tree estimation calculated as BSD between the normalized simulated species tree and the normalized estimated tree, the X axis the number of genes included in the analysis. Each boxplot includes 10 independent replicates. Different colors represent different methods (see legend). a) 1*N*_*e*_ tree height and balanced tree. b) 1*N*_*e*_ tree height and unbalanced tree. c) 10*N*_*e*_ tree height and balanced tree. d) 10*N*_*e*_ tree height and unbalanced tree. PoMo performs much better than STEM+UPGMA, and is slightly more accurate than the two concatenation approaches.

Concatenation methods have acceptable accuracy in some scenarios, but show large errors with short trees and in the “recent radiation” scenario (Figs. S11, S12). Most of these large errors do not seem to decrease when adding more data. Yet, when we use the consensus from multiple samples of the same species instead of a random sample, we notice often a small but consistent reduction in error. Accuracy of Bayesian (MrBayes) and maximum likelihood (HyPhy) methods was very similar, and overall, they both have worse or similar accuracy to PoMo.

The accuracy of STEM was generally less predictable (Figs. 4, 5). STEM seems to only provide an advantage with respect to concatenation in the “recent radiation” scenario, while it has larger error in all other scenarios. Also, the error in STEM does not seem to decrease noticeably as more genes are included into the analysis (as already observed by Leaché and Rannala 2011, with a different accuracy measure). These problems might be attributable to the particular simulation setting, with insufficient phylogenetic signal to estimate gene trees accurately, or it might be related to the particular methods used for gene tree inference. In fact, using UPGMA for gene tree estimation we obtained better accuracy than using Neighbor Joining (Figs. S13 and S14), suggesting that gene tree estimation has a large impact on the performance of STEM. We cannot exclude the possibility that with Maximum Likelihood or Bayesian gene tree inference STEM would provide better estimates, but the analysis would surely be more computationally demanding. We noticed that often STEM species trees estimates have a few branches of length 0 in scenarios with many genes and short trees. This phenomenon can happen when two gene sequences from different species are identical, and could be solved by using a Bayesian method for gene tree inference. Comparing tree estimation accuracy with the Symmetric Difference (or Robinson-Foulds metric, see Robinson and Foulds 1981) instead of BSD gave comparable results (see Figs. S15 and S16)

BEST, as noted in the previous section, is the most computationally demanding approach, and for this reason we could only run it on 4 species, and up to 10 genes (for 10 samples) or 20 genes (for 3 samples per species). On the smaller datasets we tested, BEST showed variable performance, with accuracy sometimes worse, sometimes better, but overall comparable with PoMo and most other methods (Figs. 4 and S9).

*BEAST has an underlying model similar to BEST, and shares many of its features. As mentioned earlier, running *BEAST requires increased numbers of MCMC steps to reach convergence as more genes and samples are added. It is therefore not surprising that, keeping the number of MCMC steps fixed, we do not necessarily observe an accuracy improvement in *BEAST (Figs. S17 and S18), and generally we did not reach acceptable effective sample sizes (data not shown). Overall, accuracy results for *BEAST with few genes were comparable to PoMo and BEST.

SNAPP is different from the other approaches considered so far, in that it has been proposed to study speciation events at short evolutionary times. This is particularly evident when SNAPP is applied to long trees: errors in those cases are often much higher than all other methods (Figs. S17 and S18). This could be attributed to the insufficient number of MCMC steps performed, although it seems that SNAPP tends to converge to trees with long outgroup branches (data not shown). Better explanations for this problem could therefore be that the violation of the model assumptions (biallelic unlinked sites) is causing biases, or that the prior is playing an important rule due to the small size of the datasets.

### Performance of PoMo without Population Data

Information on within-species variation is usually important to accurately infer species trees. The availability of multiple sequences from the same species can in fact be determinant to correctly inferring speciation times (see e.g. Heled and Drummond 2010), as it helps to determine the amount between-species genetic differences that are attributable to within-species variation. Therefore, a scenario in which a single sample per species is available can be particularly problematic. In PoMo, when a single sample per species is provided, we require the user to specify an input value of *θ* = 4*N*_*e*_*μ*, the degree of within-species variability. To test the performance of PoMo in this case, and its robustness to the specification of *θ*, we run PoMo on the 4-species scenarios described earlier, with a single sample per species and with various *θ* values as inputs.

Given the correct *θ* value, PoMo consistently outperforms concatenation and STEM. However, if very large error in *θ* is introduced, PoMo might give worse estimates than concatenation and STEM (Fig. 6). In general, we observe different trends for long and short trees. Long trees seem to be very robust to a wrong specification of *θ* and only small errors are observed. If the given *θ* is too small, the estimates of PoMo remain very accurate for all scenarios but the anomalous one. Short trees are more sensitive to a mis-specification of *θ* and also show an error increase for underestimated *θ* values.

**Figure 6:**
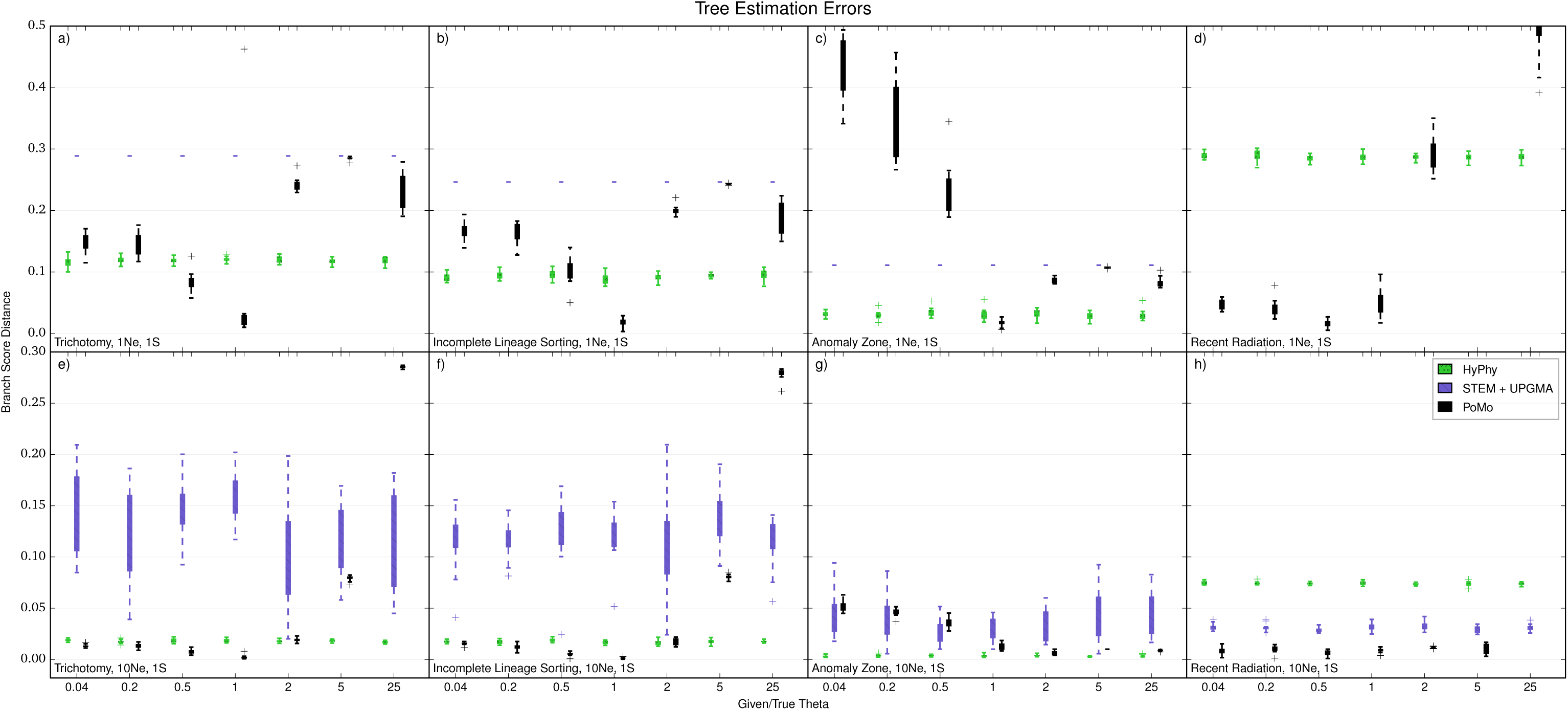
Errors in Tree Estimation with 1 Sample per Species, 4-Species-Trees. When using 1 sample per species, a value for the within-species variability _ has to be speciffed by the user. The Y axis is the error in species tree estimation calculated as BSD between the normalized simulated species tree and the normalized estimated tree. The X axis is the ratio of the guessed input _ with the simulated one. Each boxplot includes 10 independent replicates. Different colors represent different methods (see legend). a) 1*N*_*e*_ tree height and scenario with trichotomy. b) 1*N*_*e*_ tree height and ILS scenario. c) 1*N*_*e*_ tree height and anomalous species tree. d) 1*N*_*e*_ tree height and recent population radiation. e) 10*N*_*e*_ tree height and trichotomy. f) 10*N*_*e*_ tree height and ILS. g) 10*N*_*e*_ tree height and anomalous species tree. h) 10*N*_*e*_ tree height and recent population radiation. The PoMo estimates are depending on the quality of the guess for θ. We therefore do not recommend to use PoMo in this situation.

For these reasons, we suggest being cautious when using PoMo on data that only includes single samples per species, because estimates can only be trusted when *θ* is known with good confidence or when the scenario is robust to a possible mis-specification of *θ*. Yet, we expect that with the constant advancement in sequencing technologies, data sets with more than one sample per species will become increasingly predominant (see Bentley 2006).

### Application to Great Ape Data Set

Both PoMo and concatenation were run using HyPhy on the genome-wide great ape data described in *Methods*. We expect variation between consecutive runs due to different seeds and because of the random selection of samples. To assess the variability in parameter estimates, each analysis was repeated 10 times.

We can see that PoMo always infers the *Western-Lowland* and *Cross-River* gorillas to be more closely related to each other than to the *East-Lowland* gorillas (Fig. 7). Also, PoMo always infers the *Central* and *Eastern* Chimpanzee to form a clade. Both these conclusions are supported by an array of methods (Prado-Martinez et al. 2013), and also by the geographical distribution of the species. Yet, we see that concatenation places uncertainty in these observations (6 in 10 concatenation runs confirm the *Western-Lowland* and *Cross-River* gorilla clade, and 4 in 10 runs the *Central* and *Eastern* chimpanzee clade).

**Figure 7:**
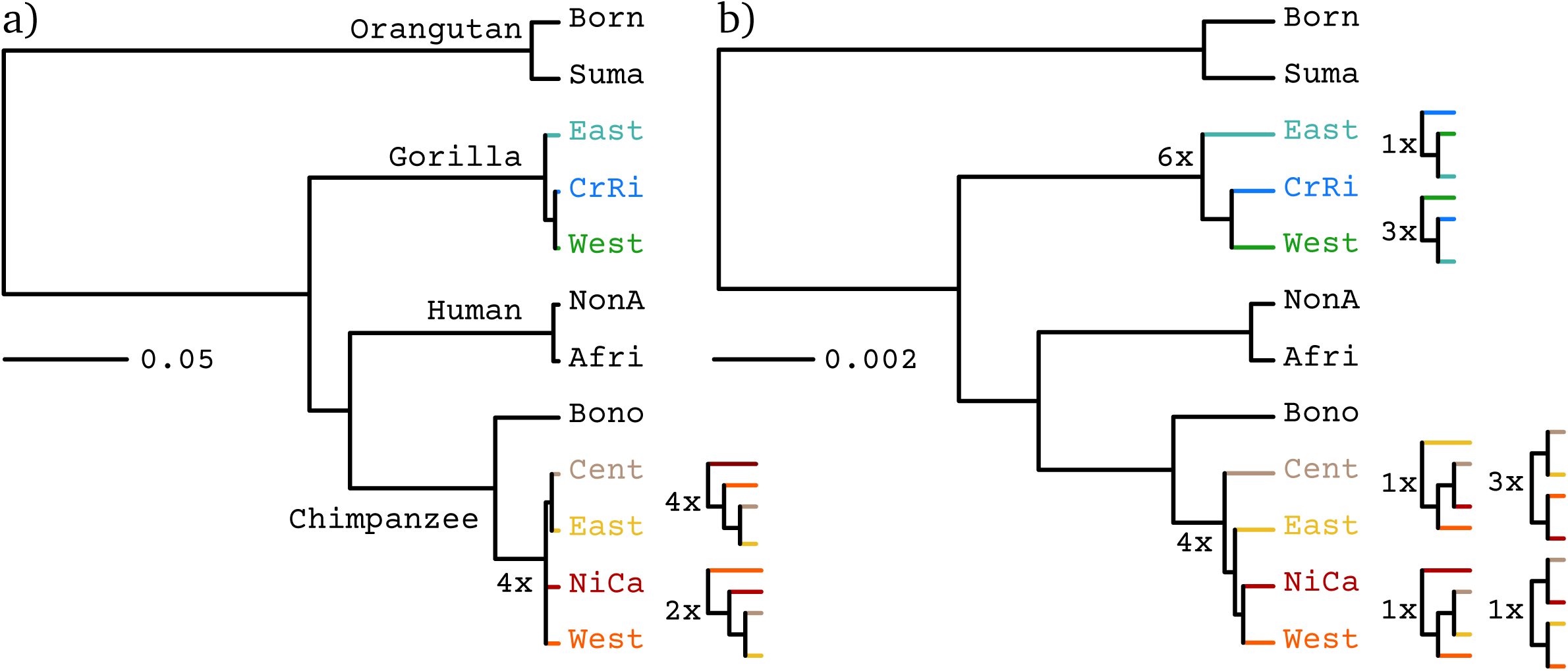
Species Tree Estimation on the Great Ape Dataset. Phylogenies were inferred using (a) PoMo and (b) concatenation. Population names are abbreviated (Born: *Bornean*, Suma: *Sumatran*, East: *Eastern*, CrRi: *Cross-River*, West: *Western*, NonA: *Non-African*, Afri: *African*, Bono: *Bonobos*, Cent: *Central*, NiCa: *Nigeria-Cameron*). The numbers indicate the abundance of the different clade topologies among different runs (we performed a total of 10 runs per method). The PoMo trees are topologically more stable than the trees estimated from the concatinated data of one randomly chosen individual per species. Interpretation of phylogenetic scales differs between the two methods. In fact, state changes in concatenation represent substitutions, while in PoMo they represent mutation and drift.

Furthermore, we can see that branches representing recent population and species splits are inferred to be more recent by PoMo than by concatenation (Fig. 7). This happens because PoMo accounts for shared and ancestral polymorphisms, while concatenation attributes all differences to divergence. Also, another reason is that concatenation interprets phylogenetic signal that is incongruent with the species tree (due to ILS) as due to multiple mutations. These patterns are relatively weak in Great Apes due to their small effective population size, but in species with higher *θ* we expect these trends to be even more pronounced. In addition to the robustness of tree topology and branch lengths we also observe that the likelihood is less variable for PoMo than for concatenation (Table S2). The average runtime of PoMo was 21 hours on a standard desktop PC (processor: Intel i5-3330S, 2.70GHz, 2 physical cores).

## CONCLUSIONS

In this study, we addressed the problem of estimating species trees in the presence of Incomplete Lineage Sorting (ILS). Using simulations covering different scenarios, we tested the computational efficiency and the accuracy of different methods. Most details of our simulations are close to those used in previous similar studies (in particular see Huang et al. 2010). We did not focus only on topology estimation, but rather evaluated the accuracy of methods in retrieving both topology and branch lengths. We also proposed a new approach for species tree estimation, PoMo, based on a recently introduced phylogenetic model of sequence evolution (De Maio et al. 2013).

With our simulations we suggest that methods such as *BEAST and BEST are not suited for the analysis of large datasets (see also Liu et al. 2009a). In fact, their computational demand is often already excessive with few genes, and increases considerably when more genes and samples are added. So, while these approaches are useful in small dataset, in particular to account for uncertainty in gene tree estimates, they are not generally applicable to exome-scale data and to large numbers of samples.

The fastest method tested was STEM, at least unless we account for the time required for gene tree estimation. In fact, STEM does not estimate the species tree from alignments, but from gene trees estimated by the user. Even when using very fast phylogenetic methods, gene tree estimation can be demanding if many loci and samples are included. Overall, STEM proved applicable to large datasets, but its estimates did not converge to the simulated values as we included more genes. This can be explained by the fact that STEM is not robust to errors in gene tree estimation.

In many cases with pervasive ILS, concatenation methods were reasonably accurate, converging to trees not very distant from the truth. Yet, in particular for cases of recent radiation when the polymorphisms are shared between species, concatenation had very high error.

We also tested the performance of SNAPP, a recent approach that does not specifically fit the assumptions of our simulations, but that has many features in common with PoMo. We were not able to obtain convergence to the correct trees with SNAPP using our limited computational resources.We cannot exclude the possibility that some specific assumption violation contributed to this problem, in fact our simulations were tailored in particular for maximum likelihood and heuristic methods, and not for Bayesian approaches.

Lastly, we showed that PoMo provides very accurate estimates, converging towards simulated values as more genes are included in the analysis, with reasonable computational demand. PoMo is slower than concatenation, although in some sense it can be considered itself a concatenation approach. The difference is mostly due to the use of a larger substitution matrix. Running PoMo never required more than a few hours on the datasets consider here, but we are working to make it faster by decreasing the dimension of the rate matrix or by using faster phylogenetic packages, such as RAxML (Stamatakis 2014) or IQTree (Nguyen et al. 2015). In fact, one of the advantages of PoMo is that it is easily exportable, and it is simple to extend with classical features of phylogenetic models. For example, presently PoMo users can choose between different mutation models, and it is possible to adopt a molecular clock, or site variation in mutation rates, and fixation biases. However, it does not yet include features of other methods such as rate variation between genes, and population size variation along the phylogeny.

The accuracy of PoMo was broadly comparable to most other methods when few genes were considered. Poor estimates in this situation can be attributable to insufficient signal. Yet, we also want to remark that PoMo assumes that all sites are unlinked: while we showed that this assumption of PoMo does not lead to biases when many independent loci are considered, caution is required when dealing with few genes. For example, in the extreme case of a single large non-recombining locus, PoMo might infer the gene tree to be the species tree with high confidence. For such scenarios, it is better to use models that account for within-locus linkage, in particular Bayesian models that also provide estimates of uncertainty, such as *BEAST ad BEST. Also, inference of species trees is problematic in the absence of information regarding within-species variation. We urge caution using PoMo with a single sample per species, and suggest to acquire additional samples unless good estimates of within-species genetic variation are known.

Apart from these limitations, PoMo can be applied to a wide selection of scenarios where most other methods are not suited. For example, PoMo can be used when intra-locus recombination is very strong, or equivalently, when loci are very small, which includes whole-genome alignments with high recombination. It is also applicable when haplotype information is not available (e.g. in pool sequencing, see Kofler et al. 2011). PoMo does not need alignment data to be arbitrarily split into loci, and is not encumbered by large numbers of samples or sites. As an example of the applicability of PoMo, we used it on a genome-wide data set comprising several samples (79) and taxa (12) of great apes. We extracted synonymous sites and collated them into a single alignment of ∼ 2.8 million bp. Using PoMo, in a few hours we could estimate species tree topologies which were more consistent with previous literature than using concatenation. Also, results using PoMo were more congruent across different runs.

Estimates of PoMo and concatenation also differed in branch lengths, in fact, as supported by our simulations, concatenation tends to overestimate short terminal branches. This phenomenon is probably due to concatenation interpreting differences between samples as divergence rather than within-species variation. In other words, concatenation attempts to estimate an average coalescent tree, rather than the species tree. Also, concatenation ignores the effects of recombination, and this can result in interpreting SNPs incongruent with the species tree as due to multiple mutations. These effects will be even more remarked for taxa with larger within-species variation *θ*. In conclusion, we think that PoMo will prove very useful in providing accurate species tree estimation from a great variety of datasets.

## SOFTWARE AVAILABILITY

PoMo is open source and can be downloaded at

~~~
https://github.com/pomo-dev/PoMo
~~~

.

## Acknowledgements

We are greatful to reviewers Laura Kubatko, Gergely Szöllőosi and Bryan Carstens for constructive comments. The computational results presented above have partly been achieved using the Vienna Scientific Cluster (VSC). We thank Christian Schlötterer for the insightful discussions regarding this project, and Jessica Hedge, Dilrini De Silva and Jane Charlesworth for comments on the manuscript. This work was supported by a grant from the Austrian Science Fund (FWF, P24551-B25 to CK). NDM and DS were members of the Vienna Graduate School of Population Genetics which is supported by a grant of the Austrian Science Fund (FWF, W1225-B20). NDM was partially supported by the Institute for Emerging Infections, funded by the Oxford Martin School.

**Online Supplementary Material**

## ONLINE APPENDIX 1: POMO TRANSITION MATRIX

Equation S2: 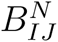, the sub-matrix describing the rates of allele frequency changes for polymorphic states with alleles *I* and *J*. The symbol 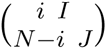 with 1 *≤ i ≤ N –* 1 represents the polymorphic state associated to alleles {*I, J*} and to frequency *i/*(*N*) of allele *I*. States representing a fixed allele are indicated by the corresponding nucleotide. Frequency change rates are defined from a Moran model with no selection:

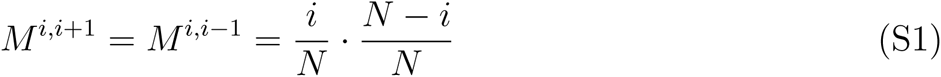

Equation S3: *Q*_*N*_, the matrix of instantaneous state change rates for PoMo*N*. *κ* is the transition rate, while *μ* is the transverion rate. For the general matrix accounting for different mutation and fixation biases (not used in this work) see De Maio, Schlötterer and Kosiol (2013).

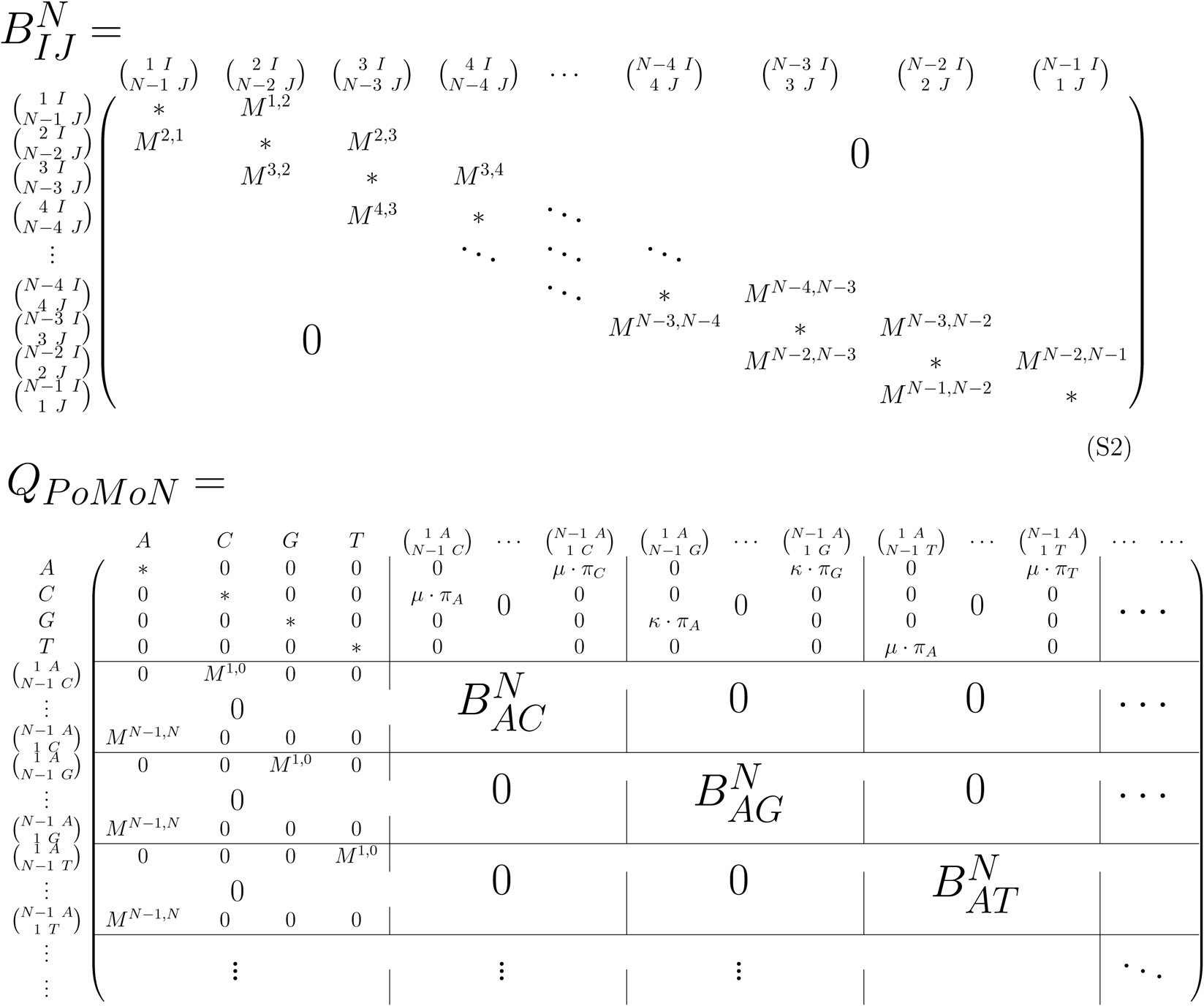

## ONLINE APPENDIX 2: SUPPLEMENTARY FIGURES

**Figure S1:**
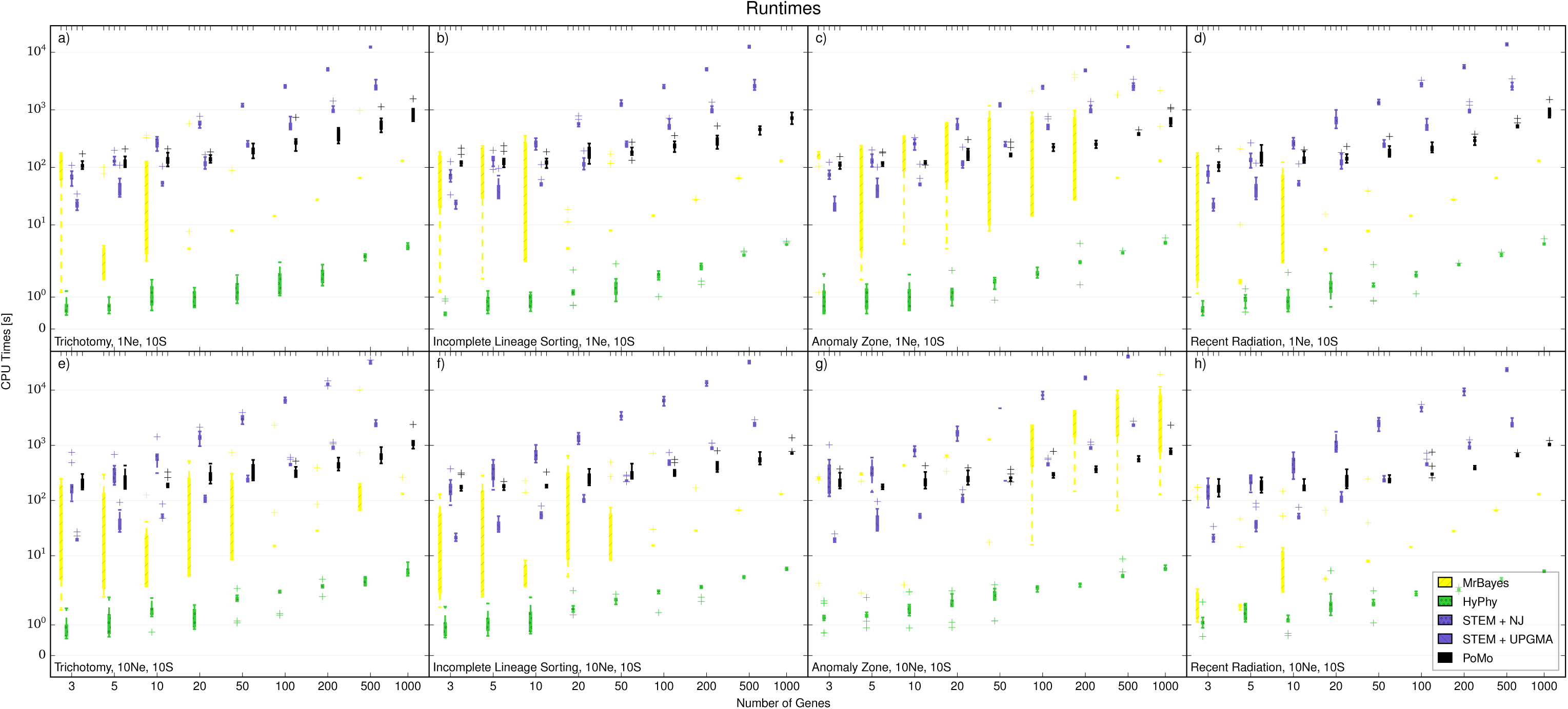
Computational Demand of STEM and Concatenation with 10 Samples per Species. On the Y axis is the computational time measured in seconds. On the X axis is the number of genes considered. Different colors represent different methods. Each boxplot. includes 10 independent replicates, a) 1*N*_*e*_ tree height and scenario with trichotomy, b) 1*N*_*e*_ tree height and ILS scenario, c) 1*N*_*e*_ tree height and anomalous species tree, d) 1*N*_*e*_ tree height and recent population radiation, e) 10*N*_*e*_ tree height and trichotomy, f) 10*N*_e_ tree height and ILS. g) 10*N*_*e*_ tree height and anomalous species tree, h) l0*N*_*e*_ tree height and recent population radiation.

**Figure S2:**
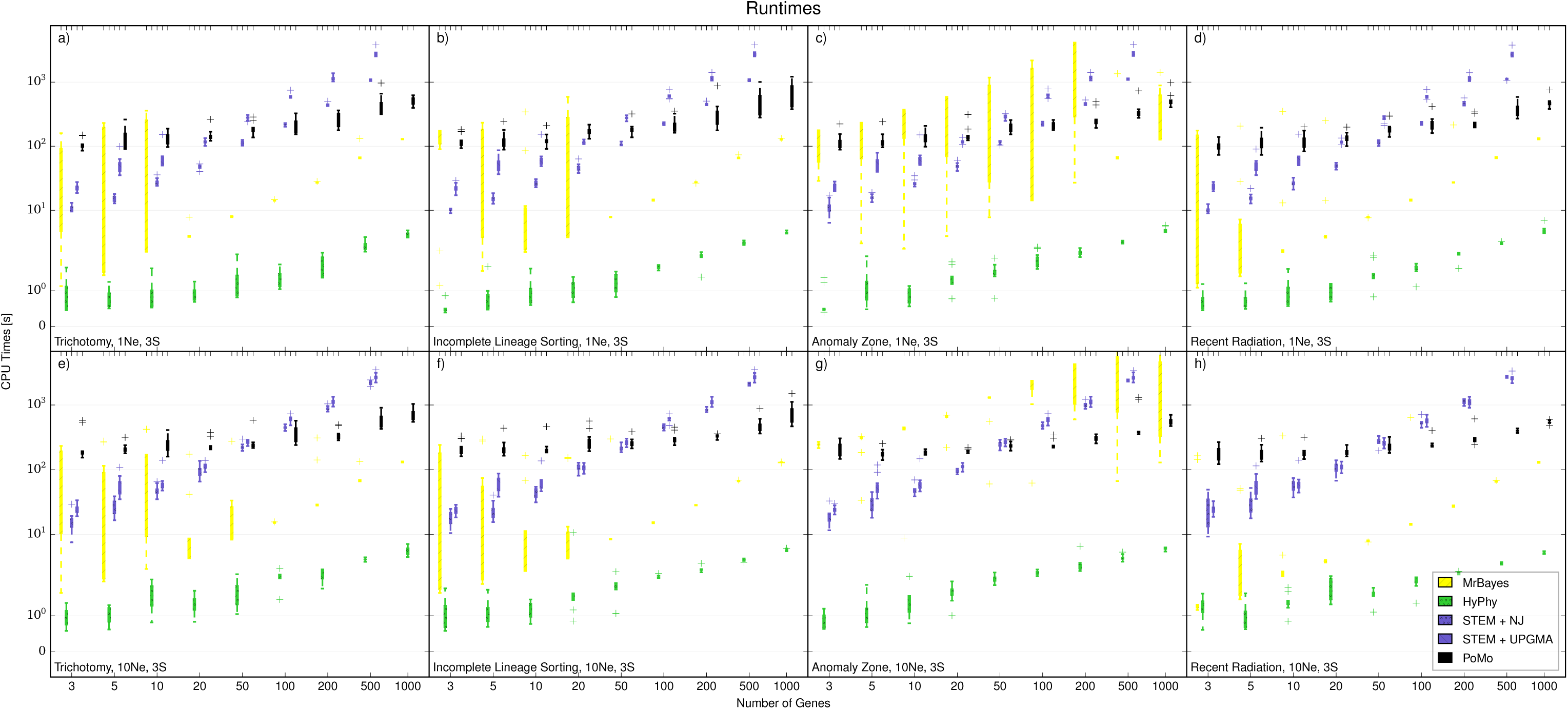
Computational Demand of STEM and Concatenation with 3 Samples per Species. On the Y axis is the computational time measured in seconds. On the X axis is the number of genes considered. Different colors represent different methods. Each boxplot includes 10 independent replicates. a) 1*N*_*e*_ tree height and scenario with trichotomy. b) 1*N*_*e*_ tree height and ILS scenario. c) 1*N*_*e*_ tree height and anomalous species tree. d) 1*N*_*e*_ tree height and recent population radiation. e) 10*N*_*e*_ tree height and trichotomy. f) 10*N*_*e*_ tree height and ILS. g) 10*N*_*e*_ tree height and anomalous species tree. h) 10*N*_*e*_ tree height and recent population radiation.

**Figure S3:**
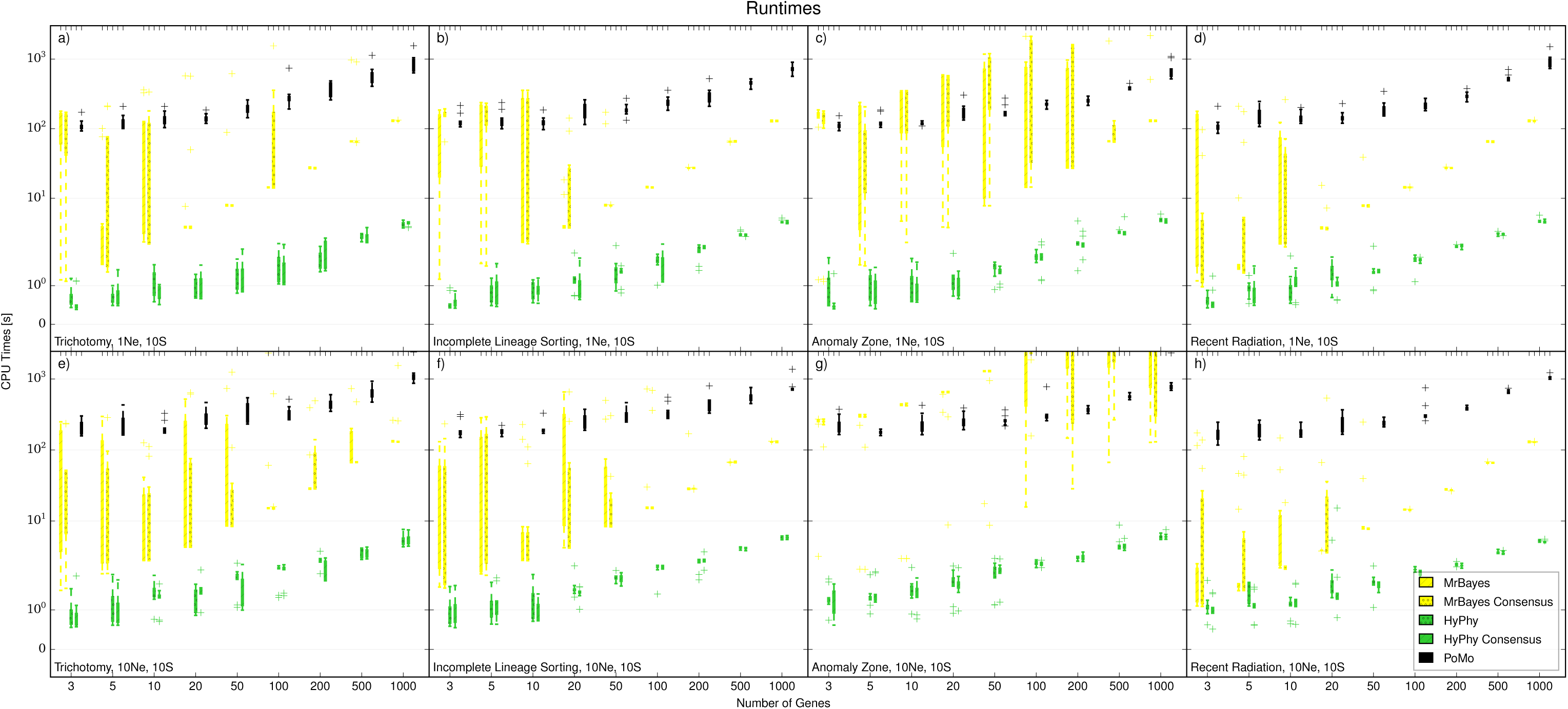
Computational Times Required by Concatenation with 10 Samples per Species. On the Y axis is the computational time measured in seconds. On the X axis is the number of genes considered. Different colors represent different methods. Each boxplot. includes 10 independent replicates, a) 1*N*_*e*_ tree height and scenario with trichotomy, b) 1*N*_*e*_ tree height and ILS scenario, c) 1*N*_*e*_ tree height and anomalous species tree, d) 1*N*_*e*_ tree height and recent population radiation, e) 10*N*_*e*_ tree height and trichotomy, f) 10*N*_*e*_ tree height and ILS. g) 10*N*_*e*_ tree height and anomalous species tree, h) l0*N*_*e*_ tree height and recent population radiation.

**Figure S4:**
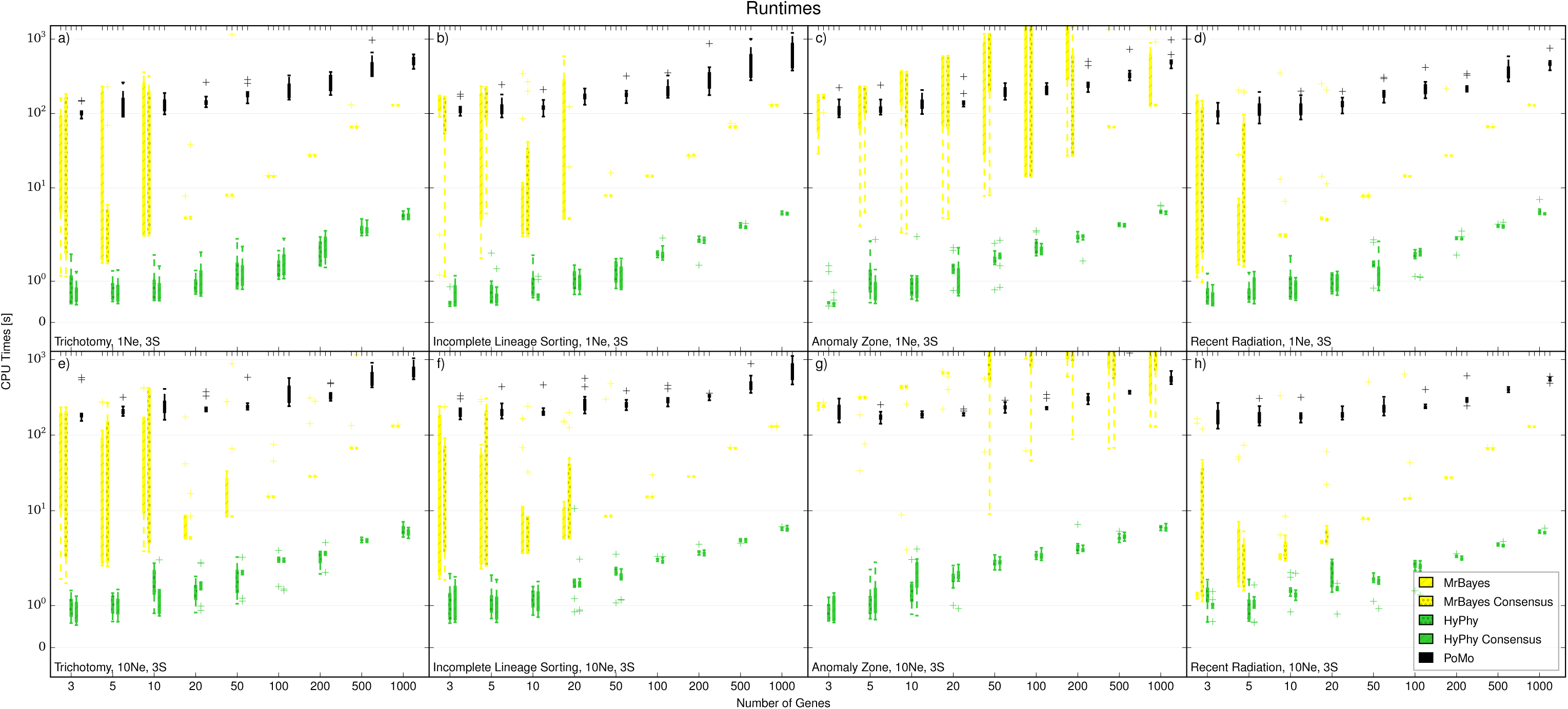
Computational Times Required by Concatenation with 3 Samples per Species. On the Y axis is the computational time measured in seconds. On the X axis is the number of genes considered. Different colors represent different methods. Each boxplot includes 10 independent replicates. a) 1*N*_*e*_ tree height and scenario with trichotomy. b) 1*N*_*e*_ tree height and ILS scenario. c) 1*N*_*e*_ tree height and anomalous species tree. d) 1*N*_*e*_ tree height and recent population radiation. e) 10*N*_*e*_ tree height and trichotomy. f) 10*N*_*e*_ tree height and ILS. g) 10*N*_*e*_ tree height and anomalous species tree. h) 10*N*_*e*_ tree height and recent population radiation

**Figure S5:**
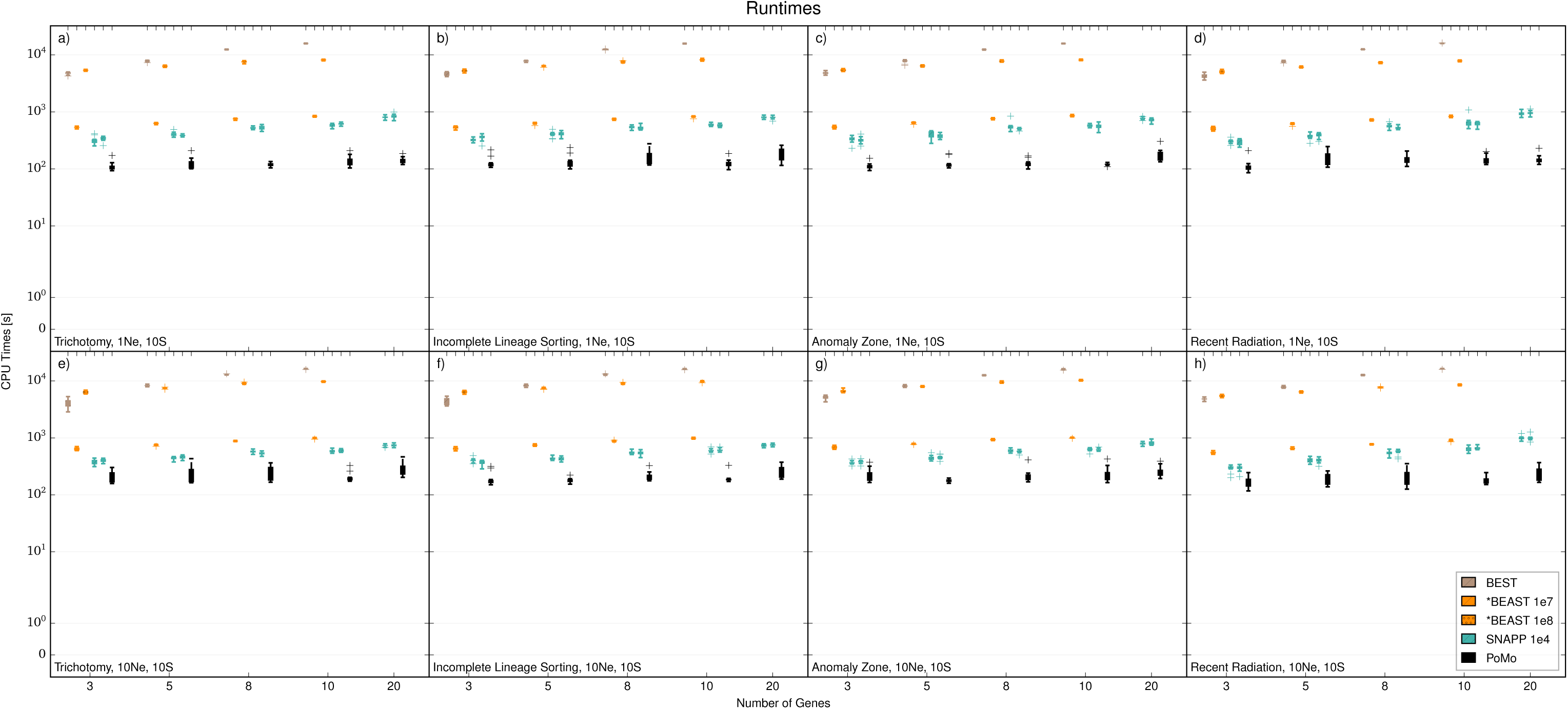
Computational Times Required by Bayesian Methods with 10 Samples per Species. On the Y axis is the computational time measured in seconds. On the X axis is the number of genes considered. Different colors represent different methods. Each boxplot. includes 10 independent replicates, a) 1*N*_*e*_ tree height and scenario with trichotomy, b) 1*N*_*e*_ tree height and ILS scenario, c) 1*N*_*e*_ tree height and anomalous species tree, d) 1*N*_*e*_ tree height and recent population radiation, e) 10*N*_*e*_ tree height and trichotomy, f) 10*N*_*e*_ tree height and ILS. g) 10*N*_*e*_ tree height and anomalous species tree, h) 10*N*_*e*_ tree height and recent population radiation. *BEAST was applied with 10^7^ MCMC steps as well as 10^8^ MCMC steps on at most 20 genes. For SNAPP 10^4^ iterations have been used.

**Figure S6:**
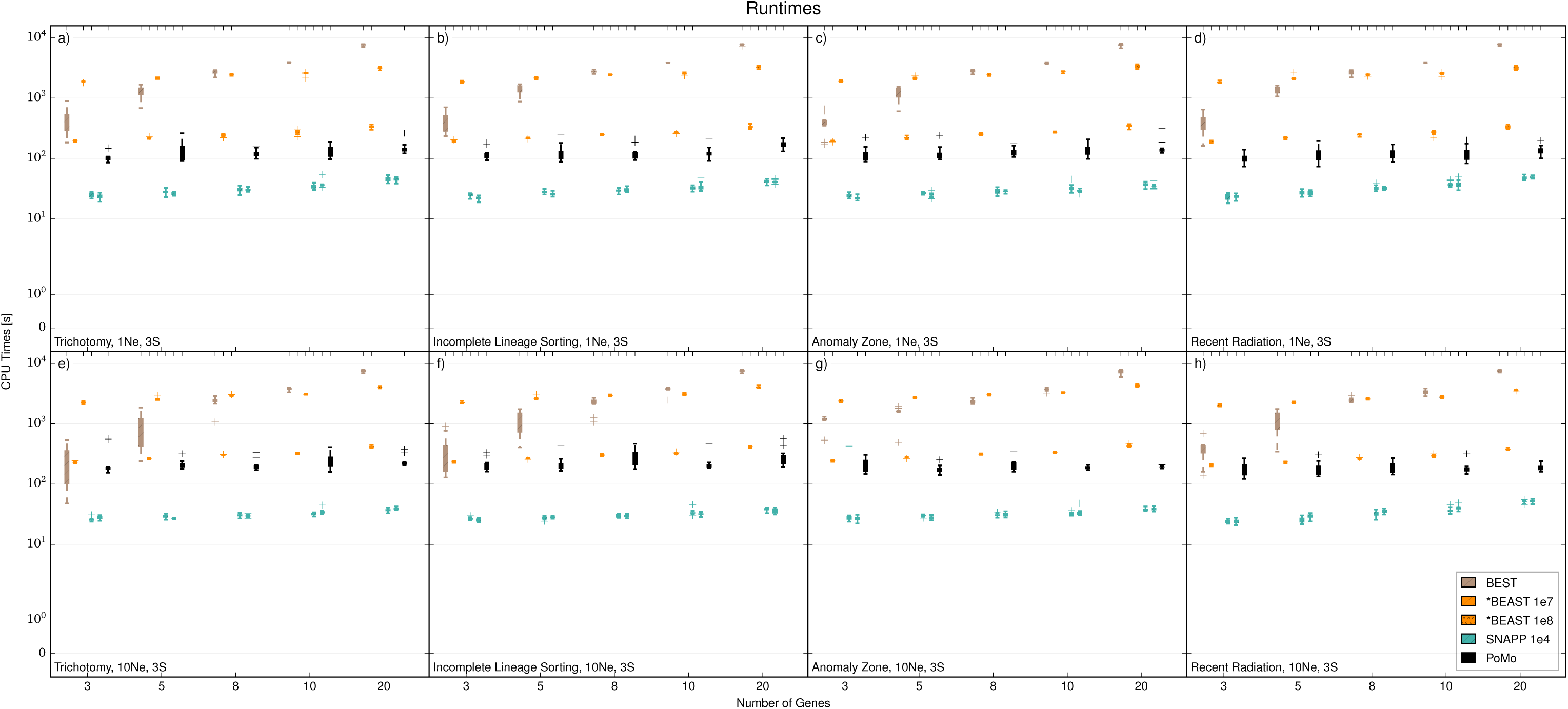
Computational Times Required by Bayesian Methods with 3 Samples per Species. On the Y axis is the computational time measured in seconds. On the X axis is the number of genes considered. Different colors represent different methods. Each boxplot. includes 10 independent replicates, a) 1*N*_*e*_ tree height and scenario with trichotomy, b) 1*N*_*e*_ tree height and ILS scenario, c) 1*N*_*e*_ tree height and anomalous species tree, d) 1*N*_*e*_ tree height and recent population radiation, e) 10*N*_*e*_ tree height and trichotomy, f) 10*N*_*e*_ tree height and ILS. g) 10*N*_*e*_ tree height and anomalous species tree, h) 10*N*_*e*_ tree height and recent population radiation. *BEAST was applied with 10^7^ MCMC steps as well as 10^8^ MCMC steps on at most 20 genes. For SNAPP 10^4^ iterations have been used.

**Figure S7:**
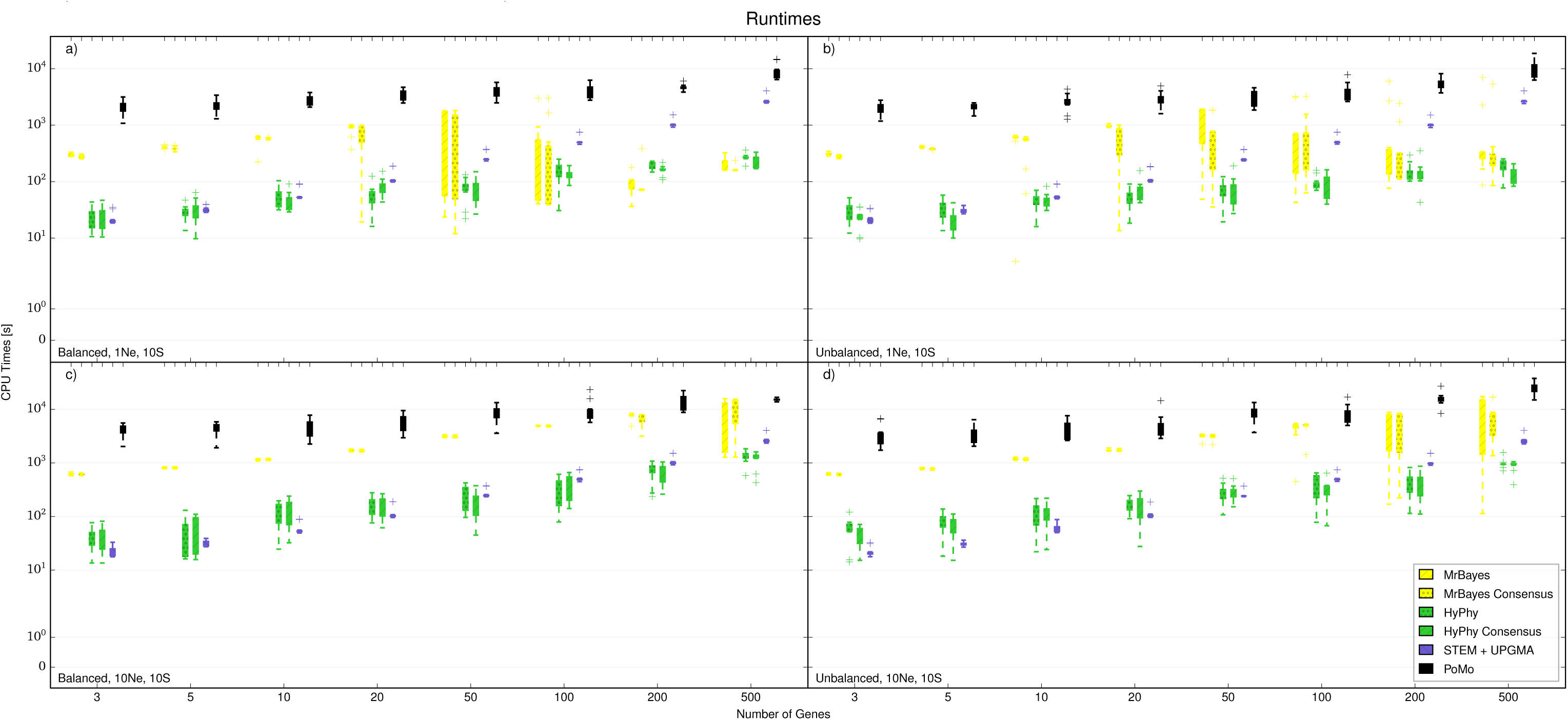
Computational Times Required with 8 Species and 10 Samples per Species. On the Y axis is the computational time measured in seconds. On the X axis is the number of genes considered. Different colors represent different methods. Each boxplot includes 10 independent replicates. a) 1*N*_*e*_ tree height and balanced tree. b) 1*N*_*e*_ tree height and unbalanced tree. c) 10*N*_*e*_ tree height and balanced tree. d) 10*N*_*e*_ tree height and unbalanced tree.

**Figure S8:**
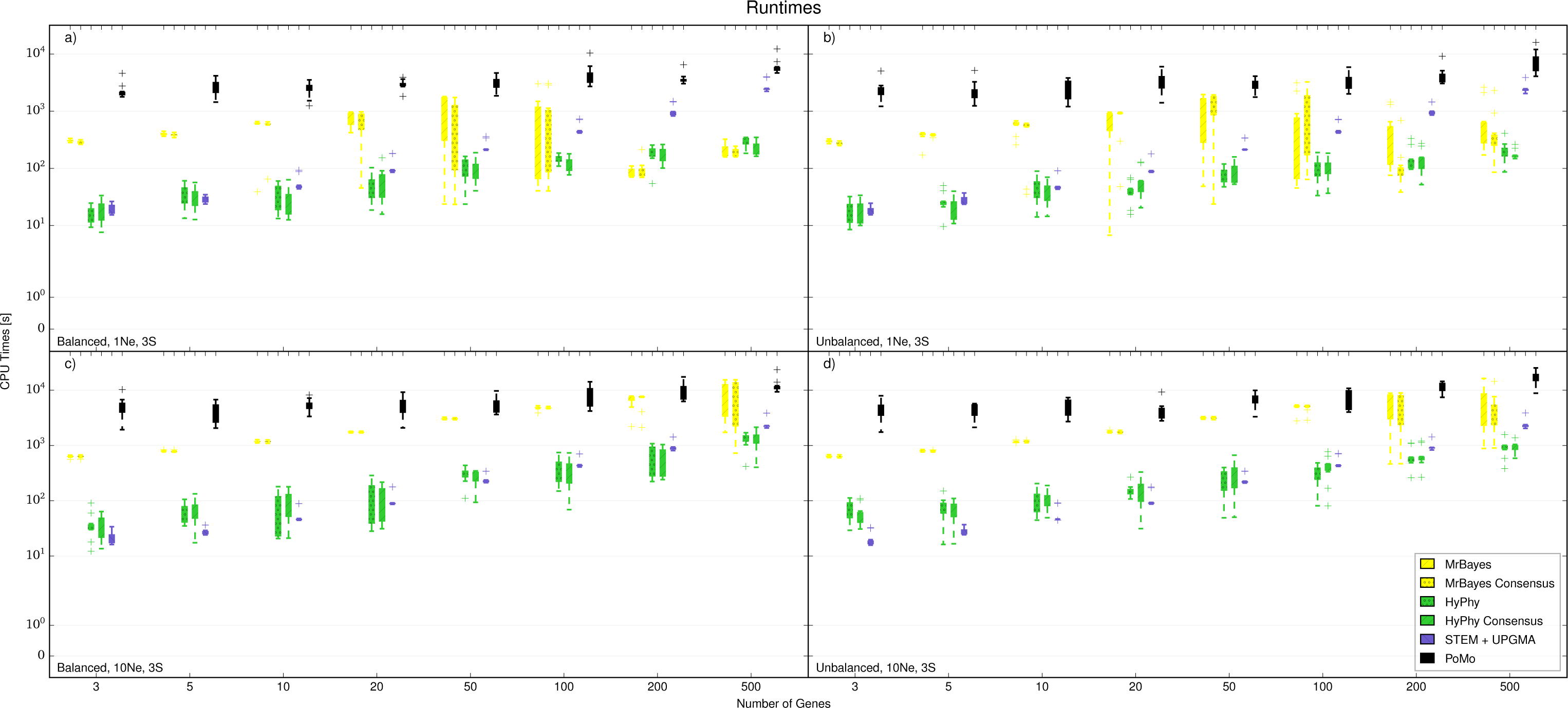
Computational Times Required with 8 Species and with 3 Samples per Species. On the Y axis is the computational time measured in seconds. On the X axis is the number of genes considered. Different colors represent different methods. Each boxplot includes 10 independent replicates. a) 1*N*_*e*_ tree height and balanced tree. b) 1*N*_*e*_ tree height and unbalanced tree. c) 10*N*_*e*_ tree height and balanced tree. d) 10*N*_*e*_ tree height and unbalanced tree.

**Figure S9:**
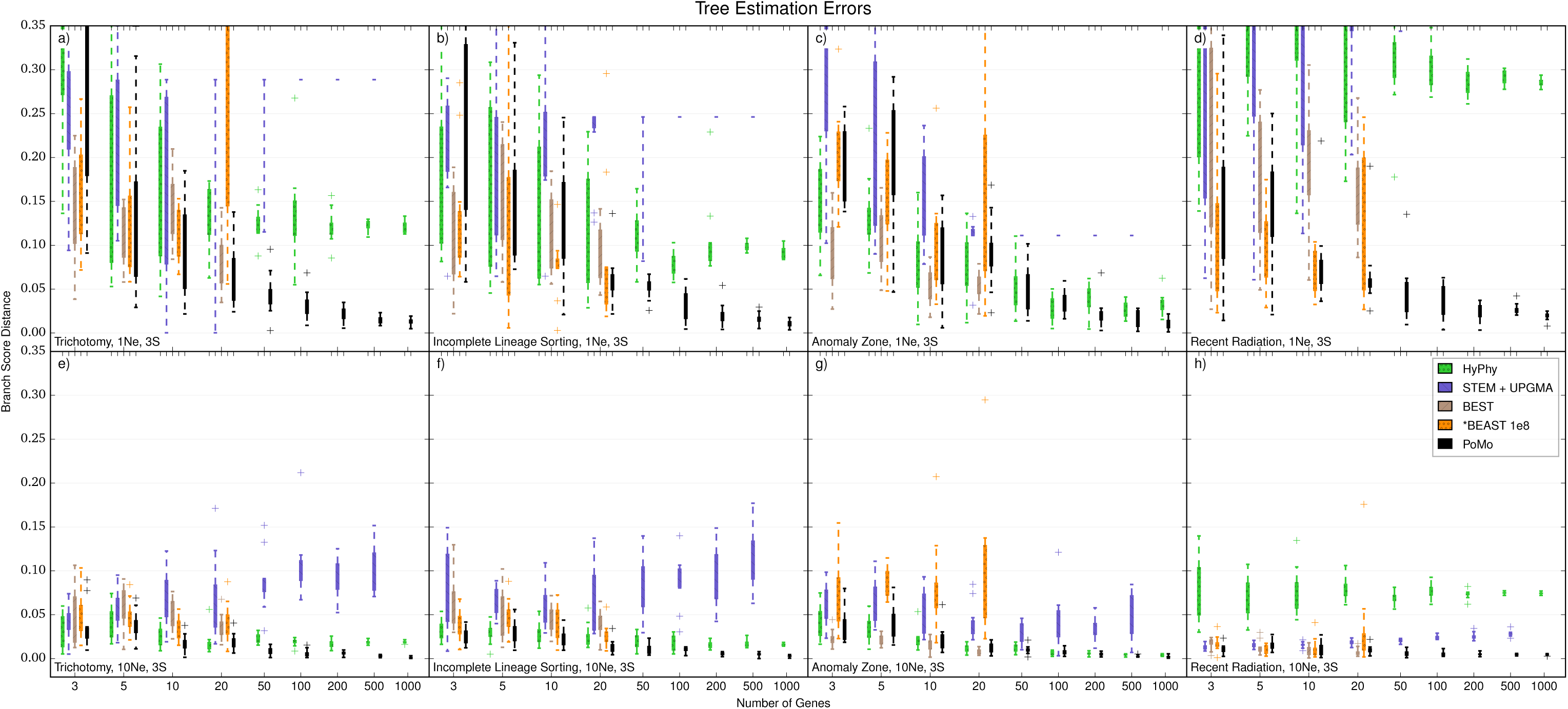
Errors in Tree Estimation with 3 Samples per Species. On the Y axis is the error in species tree estimation calculated as the Branch Score Distance (BSD) between the simulated species tree and the estimated tree. All trees are normalized prior to comparison. Each boxplot includes 10 independent replicates. On the X axis is the number of genes simulated for each replicate. Different colors represent different methods. a) 1*N*_*e*_ tree height and scenario with trichotomy. b) 1*N*_*e*_ tree height and ILS scenario. c) 1*N*_*e*_ tree height and anomalous species tree. d) 1*N*_*e*_ tree height and recent population radiation. e) 10*N*_*e*_ tree height and trichotomy. f) 10*N*_*e*_ tree height and ILS. g) 10*N*_*e*_ tree height and anomalous species tree. h) 10*N*_*e*_ tree height and recent population radiation. Some boxplot parts exceed the maximum plotted errors.

**Figure S10:**
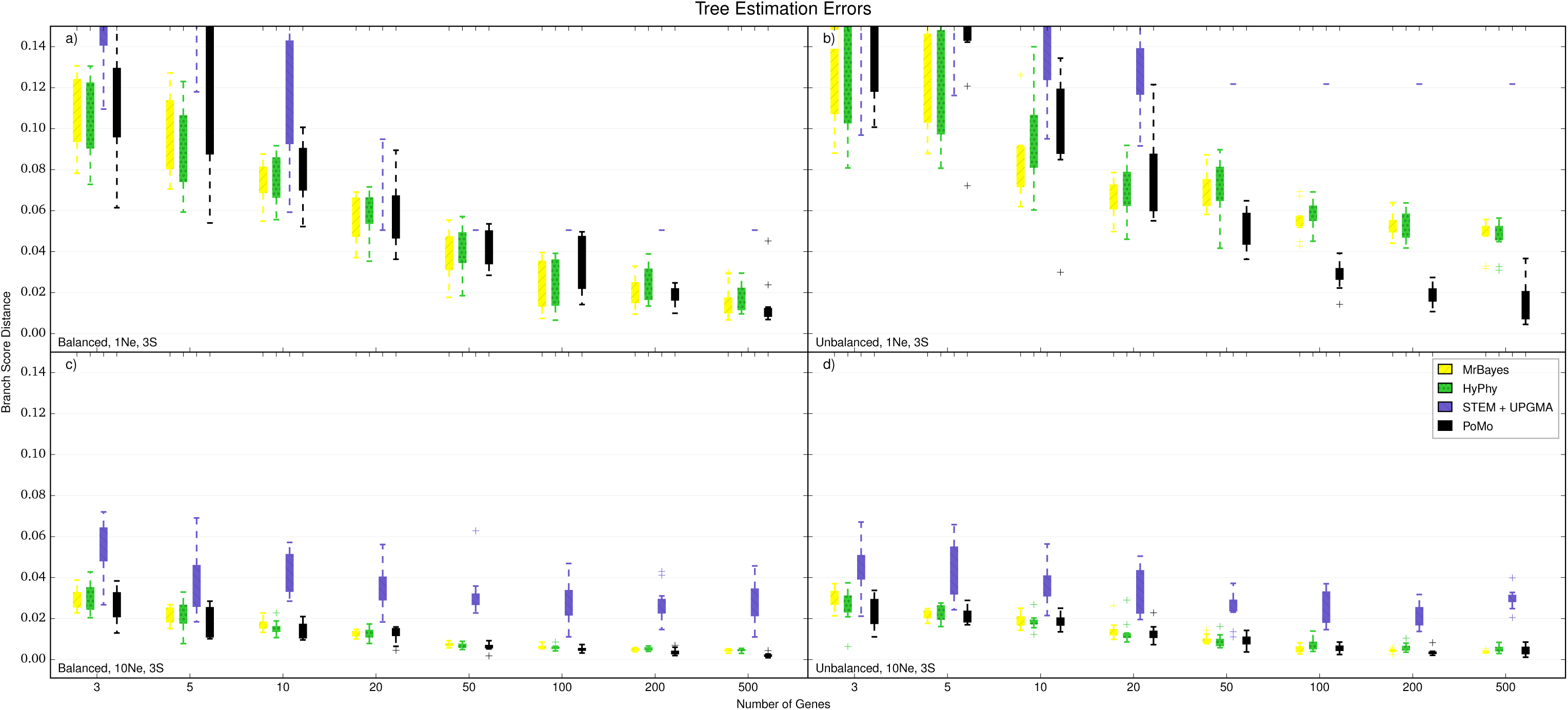
Errors in Tree Estimation with 8 Species and 3 Samples per Species. On the Y axis is the error in species tree estimation calculated as the Branch Score Distance (BSD) between the simulated species tree and the estimated tree. All trees are normalized prior to comparison. Each boxplot includes 10 independent replicates. On the X axis is the number of genes simulated for each replicate. Different colors represent different methods, a) 1*N*_*e*_ tree height and balanced tree, b) 1*N*_*e*_ tree height and unbalanced tree, c) l0*N*_*e*_ tree height and balanced tree, d) 10*N*_*e*_ tree height and unbalanced tree. Some boxplot parts exceed the maximum plotted errors.

**Figure S11:**
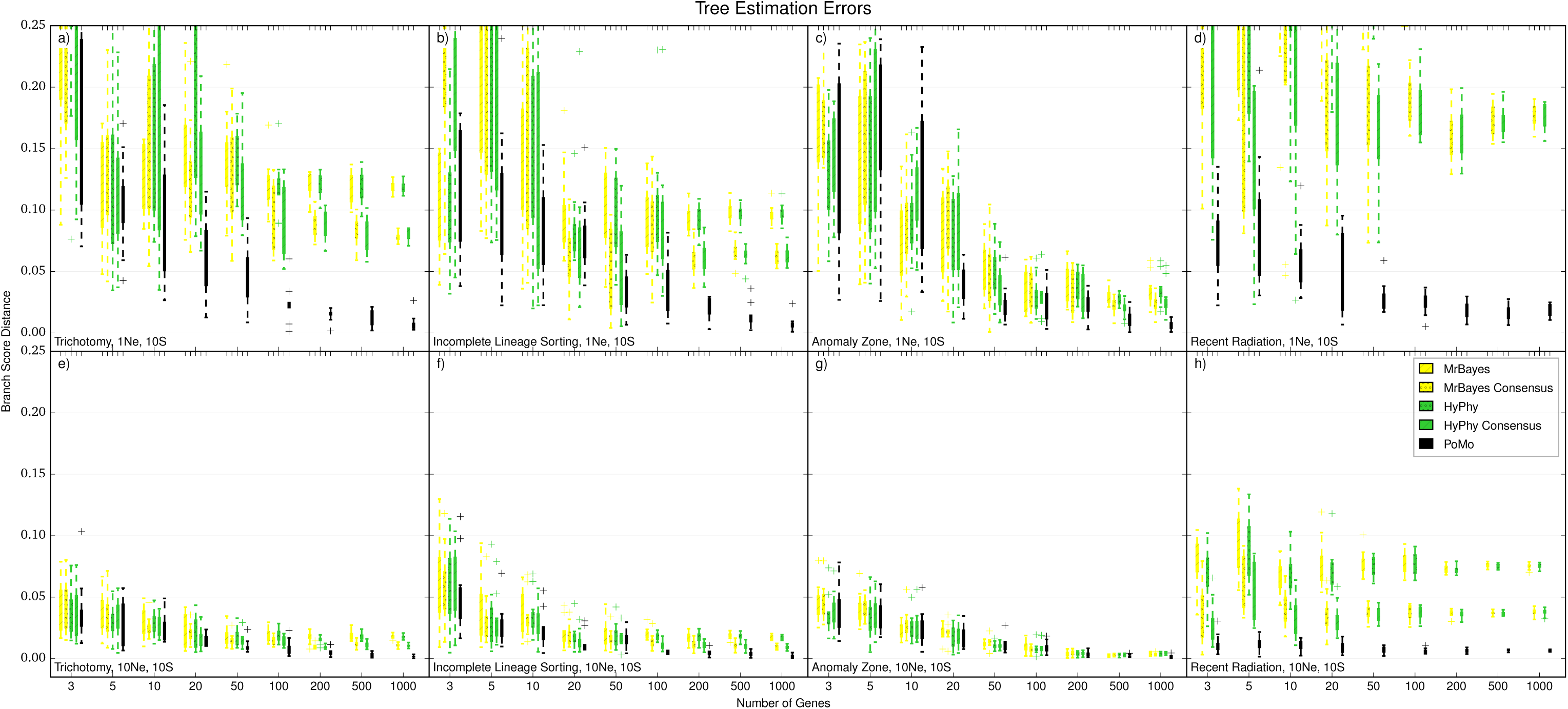
Errors in Tree Estimation of Concatenation Approaches with 10 Samples per Species. On the Y axis is the error in species tree estimation calculated as the Branch Score Distance (BSD) between the simulated species tree and the estimated tree. All trees are normalized prior to comparison. Each boxplot includes 10 independent replicates. On the X axis is the number of genes simulated for each replicate. Different colors represent different methods. “Consensus” refers to the strategy of selecting the most frequent allele in population samples, instead of selecting the allele from one random sample of the population. a) 1*N*_*e*_ tree height and scenario with trichotomy. b) 1*N*_*e*_ tree height and ILS scenario. c) 1*N*_*e*_ tree height and anomalous species tree. d) 1*N*_*e*_ tree height and recent population radiation. e) 10*N*_*e*_ tree height and trichotomy. f) 10*N*_*e*_ tree height and ILS. g) 10*N*_*e*_ tree height and anomalous species tree. h) 10*N*_*e*_ tree height and recent population radiation. Some boxplot parts exceed the maximum plotted errors.

**Figure S12:**
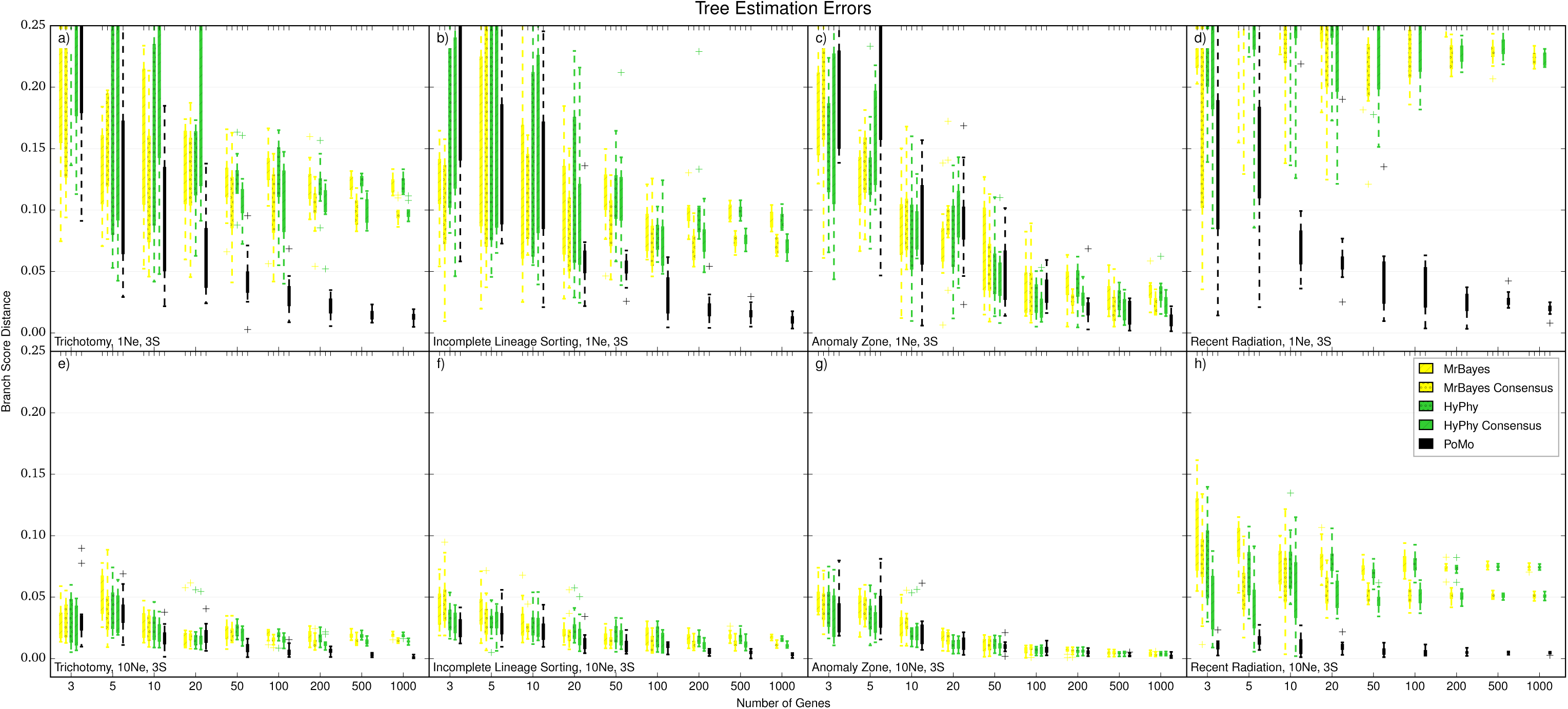
Errors in Tree Estimation of Concatenation Approaches with 3 Samples per Species. On the Y axis is the error in species tree estimation calculated as the Branch Score Distance (BSD) between the simulated species tree and the estimated tree. All trees are normalized prior to comparison. Each boxplot includes 10 independent replicates. On the X axis is the number of genes simulated for each replicate. Different colors represent different methods. “Consensus” refers to the strategy of selecting the most frequent allele in population samples, instead of selecting the allele from one random sample of the population, a) 1*N*_*e*_ tree height and scenario with trichotomy, b) 1*N*_*e*_ tree height and ILS scenario, c) 1*N*_*e*_ tree height and anomalous species tree, d) 1*N*_*e*_ tree height and recent population radiation, e) 10*N*_*e*_ tree height and trichotomy, f) 10*N*_*e*_ tree height and ILS. g) l0*N*_*e*_ tree height and anomalous species tree, h) 10*N*_*e*_ tree height and recent population radiation. Some boxplot parts exceed the maximum plotted errors.

**Figure S13:**
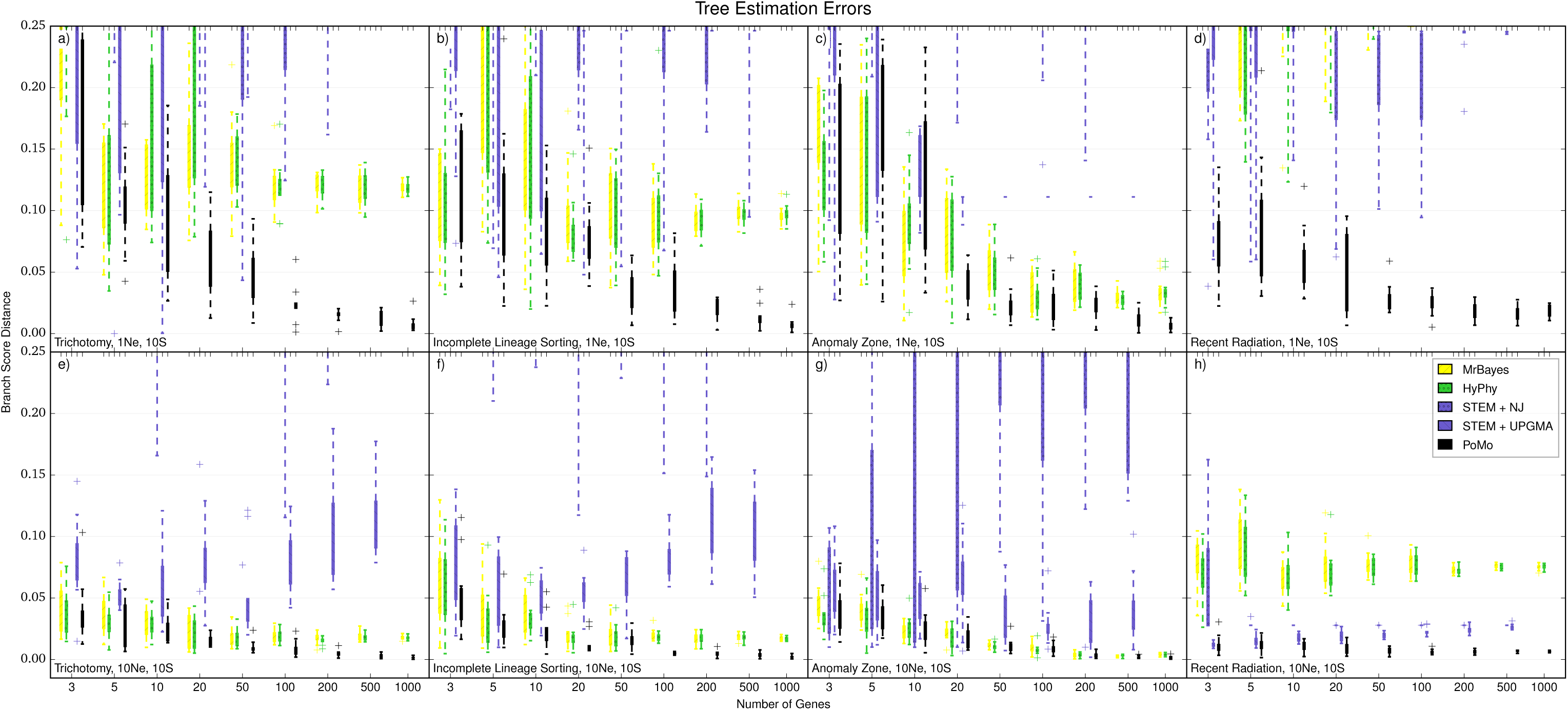
Errors in Tree Estimation of STEM and Concatenation Approaches with 10 Samples per Species. On the Y axis is the error in species tree estimation calculated as the Branch Score Distance (BSD) between the simulated species tree and the estimated tree. All trees are normalized prior to comparison. Each boxplot includes 10 independent replicates. On the X axis is the number of genes simulated for each replicate. Different colors represent different methods. a) 1*N*_*e*_ tree height and scenario with trichotomy. b) 1*N*_*e*_ tree height and ILS scenario. c) 1*N*_*e*_ tree height and anomalous species tree. d) 1*N*_*e*_ tree height and recent population radiation. e) 10*N*_*e*_ tree height and trichotomy. f) 10*N*_*e*_ tree height and ILS. g) 10*N*_*e*_ tree height and anomalous species tree. h) 10*N*_*e*_ tree height and recent population radiation. Some boxplot parts exceed the maximum plotted errors.

**Figure S14:**
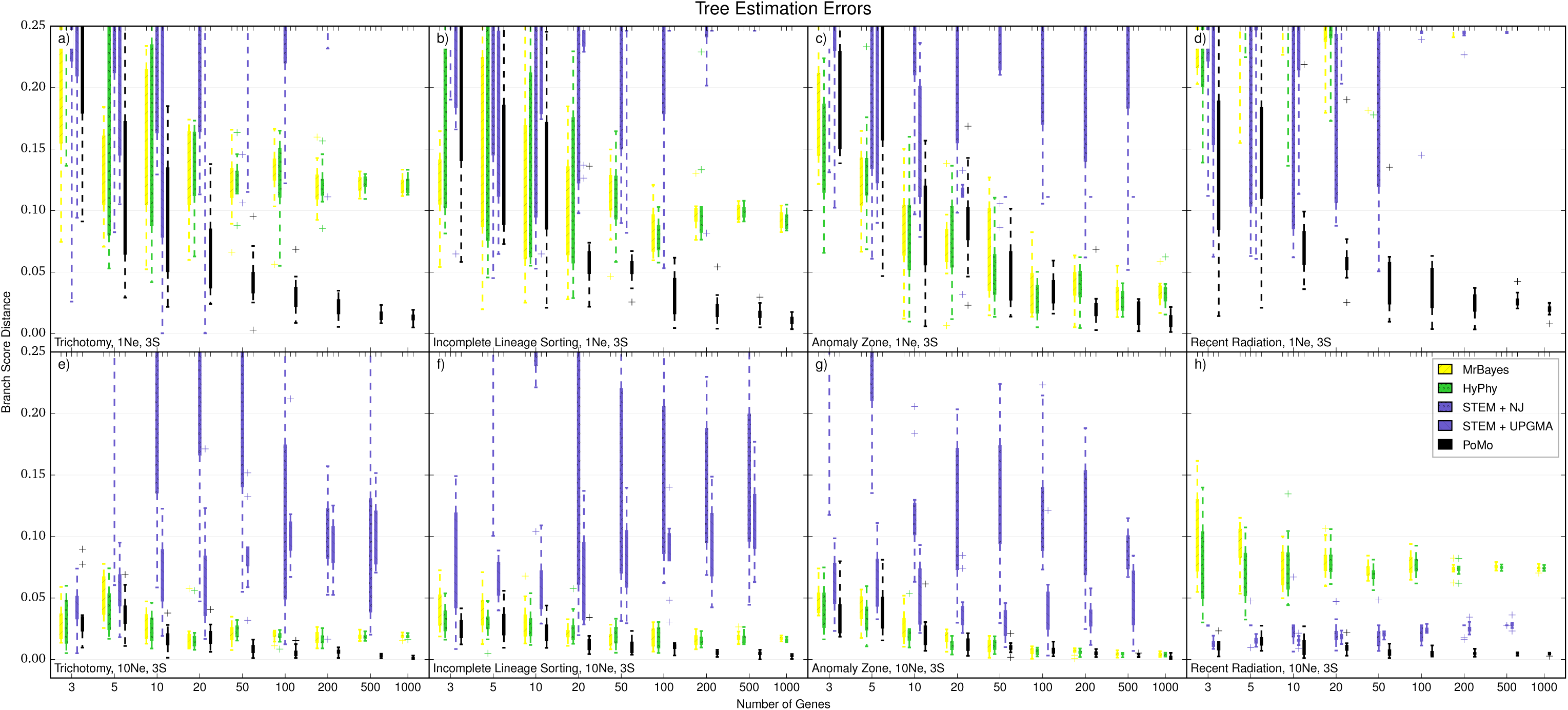
Errors in Tree Estimation of STEM and Concatenation Approaches with 3 Samples per Species. On the Y axis is the error in species tree estimation calculated as the Branch Score Distance (BSD) between the simulated species tree and the estimated tree. All trees are normalized prior to comparison. Each boxplot includes 10 independent replicates. On the X axis is the number of genes simulated for each replicate. Different colors represent different methods, a) 1*N*_*e*_ tree height and scenario with trichotomy, b) 1*N*_*e*_ tree height and ILS scenario, c) 1*N*_*e*_ tree height and anomalous species tree, d) 1*N*_*e*_ tree height and recent population radiation, e) 10*N*_*e*_ tree height and trichotomy, f) 10*N*_*e*_ tree height and ILS. g) 10*N*_*e*_ tree height and anomalous species tree, h) l0*N*_*e*_ tree height and recent population radiation. Some boxplot parts exceed the maximum plotted errors.

**Figure S15:**
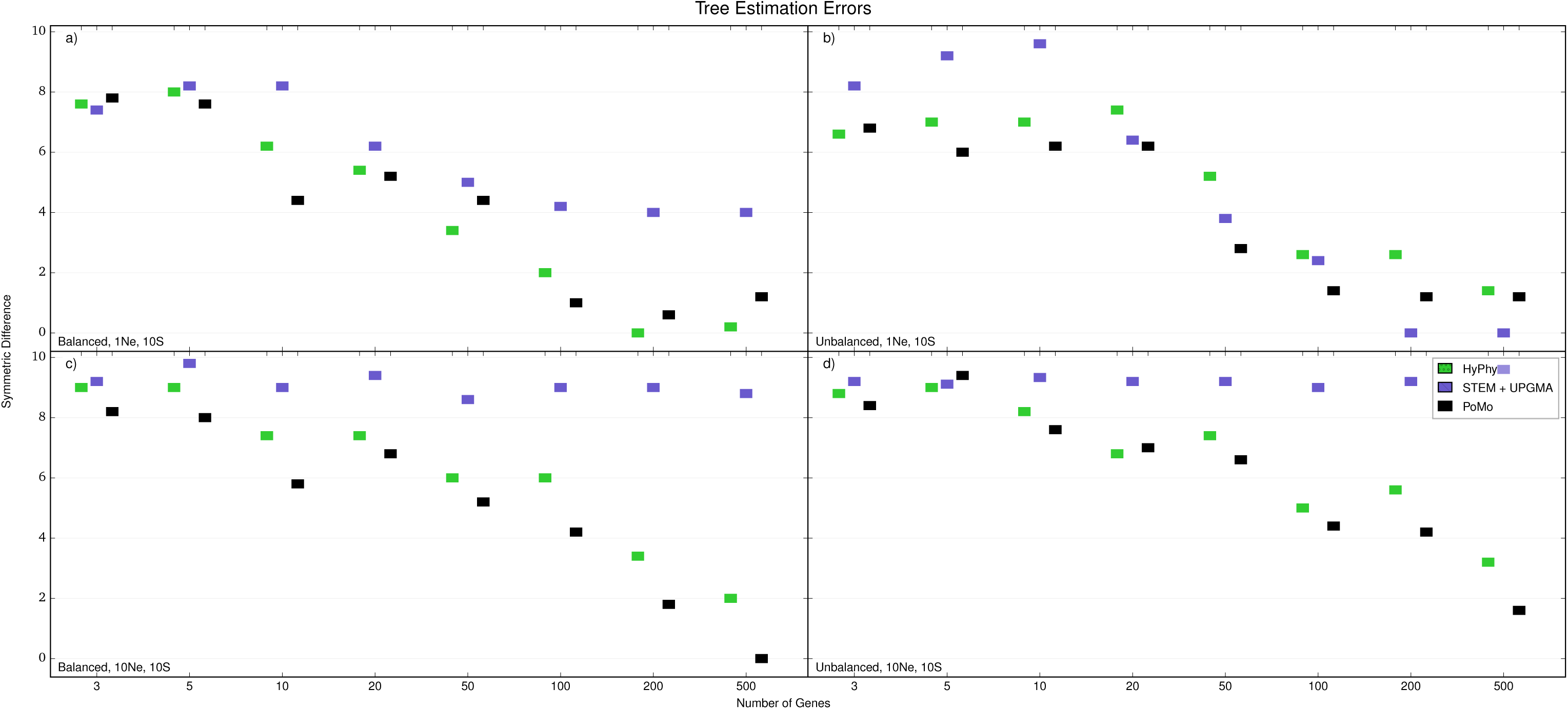
Symmetric Difference Errors in Tree Estimation with 8 Species and 10 Samples per Species. On the Y axis is the mean error in species tree estimation over the 10 replicates using the Symmetric Difference between the simulated species tree and the estimated tree. On the X axis is the number of genes simulated for each replicate. Different colors represent different methods. a) 1*N*_*e*_ tree height and balanced tree. b) 1*N*_*e*_ tree height and unbalanced tree. c) 10*N*_*e*_ tree height and balanced tree. d) 10*N*_*e*_ tree height and unbalanced tree.

**Figure S16:**
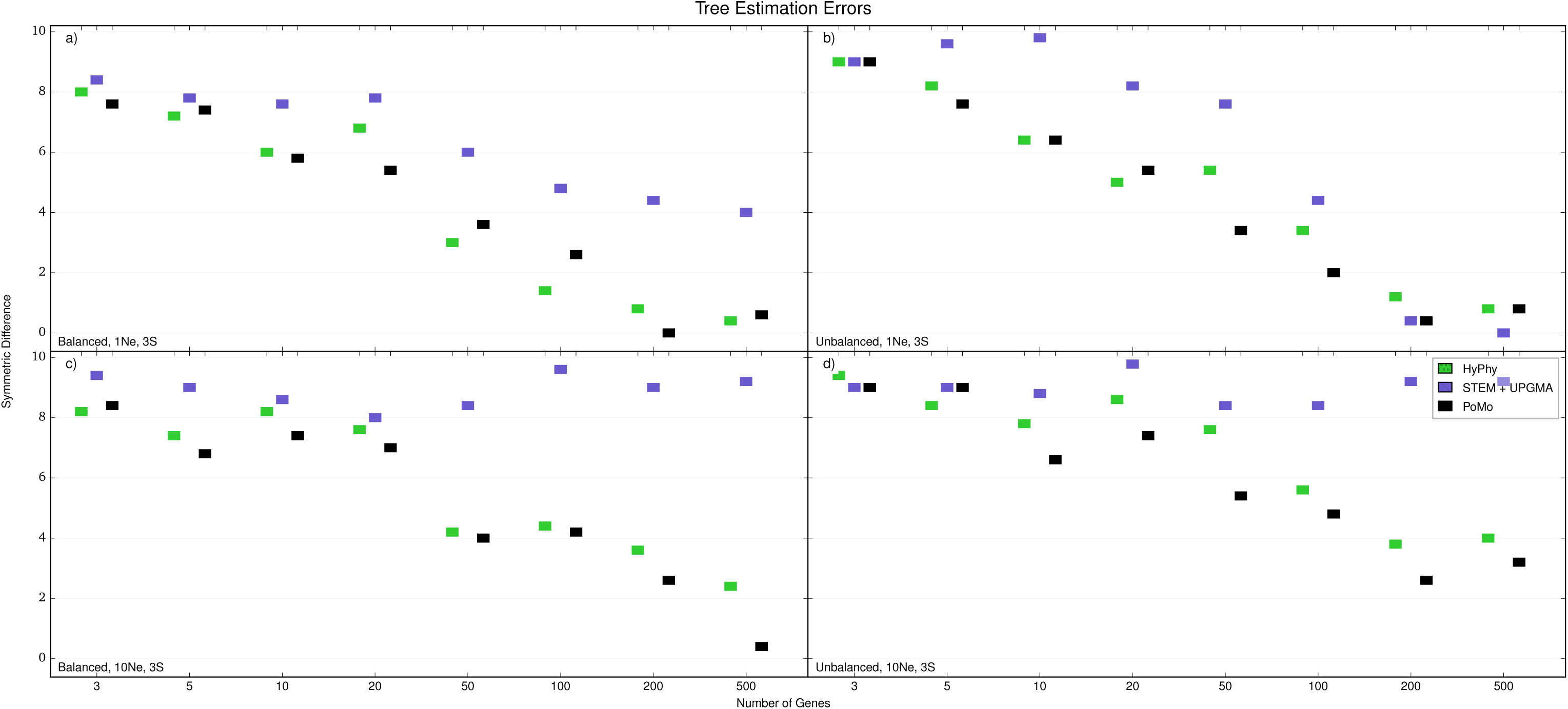
Symmetric Difference Errors in Tree Estimation with 8 Species and 10 Samples per Species. On the Y axis is the mean error in species tree estimation over the 10 replicates using the Symmetric Difference between the simulated species tree and the estimated tree. On the X axis is the number of genes simulated for each replicate. Different colors represent different methods. a) 1*N*_*e*_ tree height and balanced tree. b) 1*N*_*e*_ tree height and unbalanced tree. c) 10*N*_*e*_ tree height and balanced tree. d) 10*N*_*e*_ tree height and unbalanced tree.

**Figure S17:**
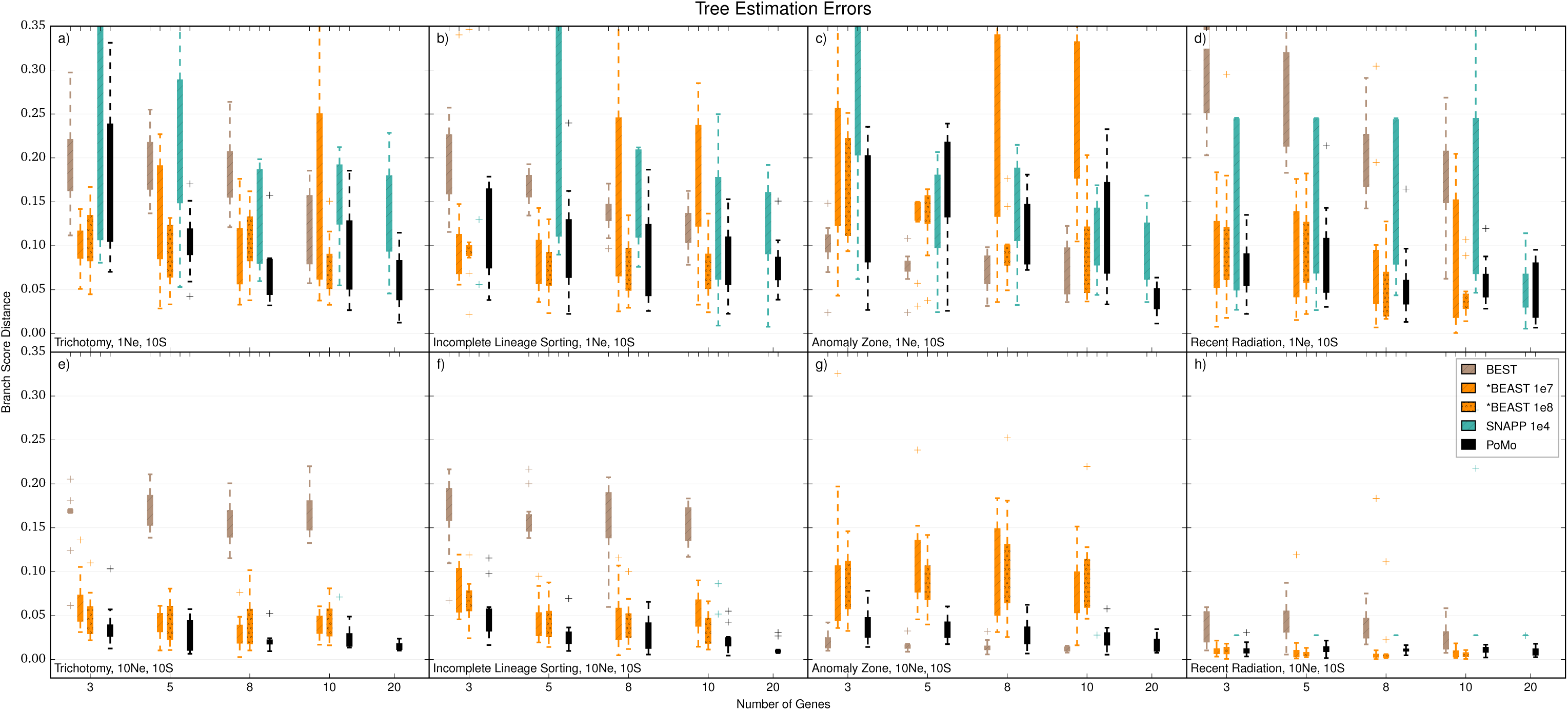
Errors in Tree Estimation of Bayesian Approaches with 10 Samples per Species. On the Y axis is the error in species tree estimation calculated as the Branch Score Distance (BSD) between the simulated species tree and the estimated tree. All trees are normalized prior to comparison. Each boxplot includes 10 independent replicates. On the X axis is the number of genes simulated for each replicate. Different colors represent different methods. “1e7” and “1e8” refer to the number of MCMC steps used (respectively 10^7^ and 10^8^). a) 1*N*_*e*_ tree height and scenario with trichotomy. b) 1*N*_*e*_ tree height and ILS scenario. c) 1*N*_*e*_ tree height and anomalous species tree. d) 1*N*_*e*_ tree height and recent population radiation. e) 10*N*_*e*_ tree height and trichotomy. f) 10*N*_*e*_ tree height and ILS. g) 10*N*_*e*_ tree height and anomalous species tree. h) 10*N*_*e*_ tree height and recent population radiation. Some boxplot parts exceed the maximum plotted errors.

**Figure S18:**
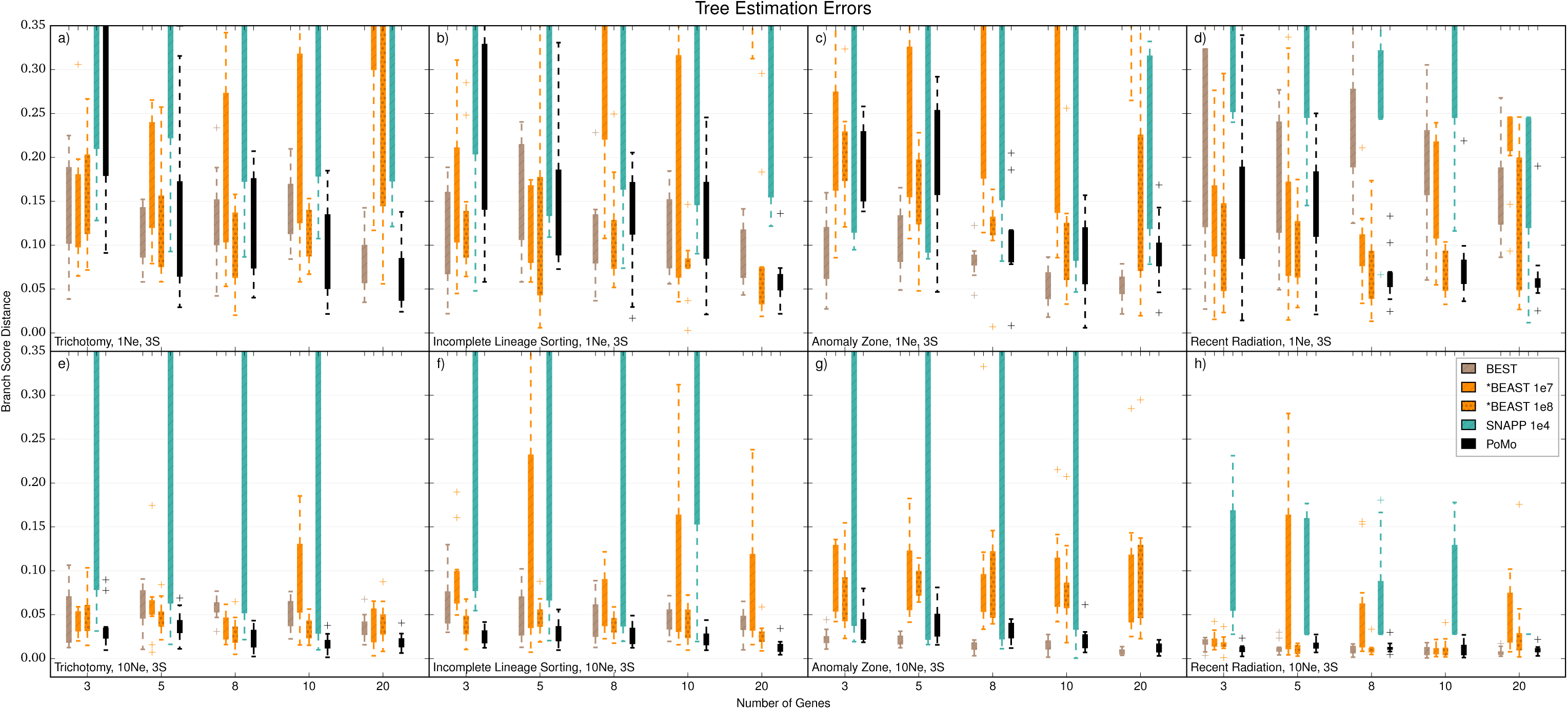
Errors in Tree Estimation of Bayesian Approaches with 3 Samples per Species. On the Y axis is the error in species tree estimation calculated as the Branch Score Distance (BSD) between the simulated species tree and the estimated tree. All trees are normalized prior to comparison. Each boxplot includes 10 independent replicates. On the X axis is the number of genes simulated for each replicate. Different colors represent different methods. “1e7” and “1e8” refer to the number of MCMC steps used (respectively 10^7^ and 10^8^). a) 1*N*_*e*_ tree height and scenario with trichotomy. b) 1*N*_*e*_ tree height and ILS scenario. c) 1*N*_*e*_ tree height and anomalous species tree. d) 1*N*_*e*_ tree height and recent population radiation. e) 10*N*_*e*_ tree height and trichotomy. f) 10*N*_*e*_ tree height and ILS. g) 10*N*_*e*_ tree height and anomalous species tree. h) 10*N*_*e*_ tree height and recent population radiation. Some boxplot parts exceed the maximum plotted errors.

## ONLINE APPENDIX 3: COMMANDS USED IN SIMULATIONS

msms for ILS, similarly for other scenarios:

*java -Xmx1g -Xmx1g -jar msms.jar* 4*sample size N genes *-t 0.01 -I 4* sample size sample size sample size sample size *-ej* T/2 *4 3 -ej* T/2+0.1 *3 2 -ej* T *2 1 -oSeqOff -T*

Where T is the total tree length (1 or 10), sample size is the number of sequences from each species (1, 3, or 10), N genes is the number of genes in the scenario.

Command for Seq-Gen:

*seq-gen -mHKY -f0.3,0.2,0.2,0.3 -t3.0 -l1000 -n1 -on -s0.0025 <* gene tree file *>* output_name

Commands for MrBayes, gene tree estimation:

*set autoclose=yes nowarn=yes execute* input file *lset nst=2 prset Brlenspr=Clock:Uniform mcmc Ngen=100000000 Samplefreq=5000 Printfreq=100000 Printall=No Diagnfreq=100000 Starttree=Parsimony Stoprule=yes Stopval=0.01 sumt conformat=simple burninfrac=0.25 Contype=Allcompat relburnin=yes quit*

Commands for STEM:

*properties:*

*run: 1*

*theta: 0.01*

*beta: 0.0005*

*mle-filename:* output_name

*species:*

*Species1: 0, 1, 2, 3, 4, 5, 6, 7, 8, 9*

*Species2: 10, 11, ….*

etc… The command line executed for STEM is:

*java -jar stem-hy.jar* settings_file

Commands for BEST 2.3:

*set autoclose=yes nowarn=yes execute* input_file;

*CHARSET G1 = 1-1000*

*CHARSET G2 = 1001-2000* etc….;

*Partion Genes =* N genes: *G1,G2,*etc…

*set partition=Genes;*

*taxset s1= 1-10;*

*taxset s2 = 11-20;*

*taxset s3= 21-30;*

*taxset s4= 31-40;*

*unlink topology=(all) brlens=(all);*

*prset best=1 brlenspr=clock:Uniform;*

*lset nst=2;*

*mcmc ngen=10000000 Stoprule = yes Stopval = 0.02 samplefreq=1000; quit*

Then, the consensus species tree is extracted:

*exe* output trees

*set autoclose=yes nowarn=yes;*

*sumt Contype=Allcompat Ntrees=1;*

*quit;*

Commands for MrBayes 3.2, species tree estimation:

*set autoclose=yes nowarn=yes execute* input_file;

*prset Shapepr=Fixed(1.0) Pinvarpr=Fixed(0.0) topologypr=uniform brlenspr=clock:uniform;*

*lset nst=2;*

*mcmc ngen=10000000 Stoprule=yes Stopval=0.001 samplefreq=500*

*Printfreq=10000 Printall=No Diagnfreq=10000;*

*sumt Conformat=Simple burninfrac=0.25 Contype=Allcompat relburnin = yes; quit;*

Commands for TreeAnnotator in BEAST 2.0.2 (for *BEAST and SNAPP output):

*-limit 0.5 -heights mean* input_file output_file

Commands for concatenation in HyPhy:

The commands are included as supplementary files. Supplementary file S1 was used to calculate a NJ tree, while file S2 was used for the NNI maximum likelihood tree.

Instructions for running PoMo:

PoMo can be downloaded at 

~~~
https://github.com/pomo-dev/PoMo
~~~

. Please follow the installation guidelines thereby.

To run PoMo, simply execute:

~~~
python path_to_PoMo/PoMo.py path_to_HYPHY/HYPHYMP path_to_data/data.txt
~~~

For the input file format, see the *sample-data* folder. Further optional arguments allow the specification of a molecular clock, different mutational models, and selection. For a full list of options run:

~~~
python path_to_PoMo/PoMo.py -h
~~~

## ONLINE APPENDIX 4: APPLICATION TO PRIMATE DATA

An exome-wide, inter- and intraspecies data set of alignments of 4-fold degenerate sites was constructed for *Homo sapiens*, *Pan troglodytes*, *Pan paniscus*, *Gorilla beringei*, *Gorilla gorilla*, *Pongo abellii* and *Pongo pygmaeus* (respectively human, chimpanzee, bonobo, eastern lowland gorilla, cross river and western lowland gorilla, Sumatran and Bornean orangutan; data from (Prado-Martinez et al. 2013)).

Multiple Consensus CoDing Sequence (CCDS) alignments that provide information on the species level (orthologous genes) were combined with the variation data on the population level from Prado-Martinez et al. (2013) to create the input data for PoMo. This was done with scripts that are provided together with PoMo. They are well documented elsewhere (http://pomo.readthedocs.org/en/v1.1.0/). The general procedure is described below.

## Interspecies Data — Consensus Coding Sequence Alignments

The CCDS alignments were downloaded from the UCSC table browser (http://genome.ucsc.edu) with the following preferences:

**Table S1.**
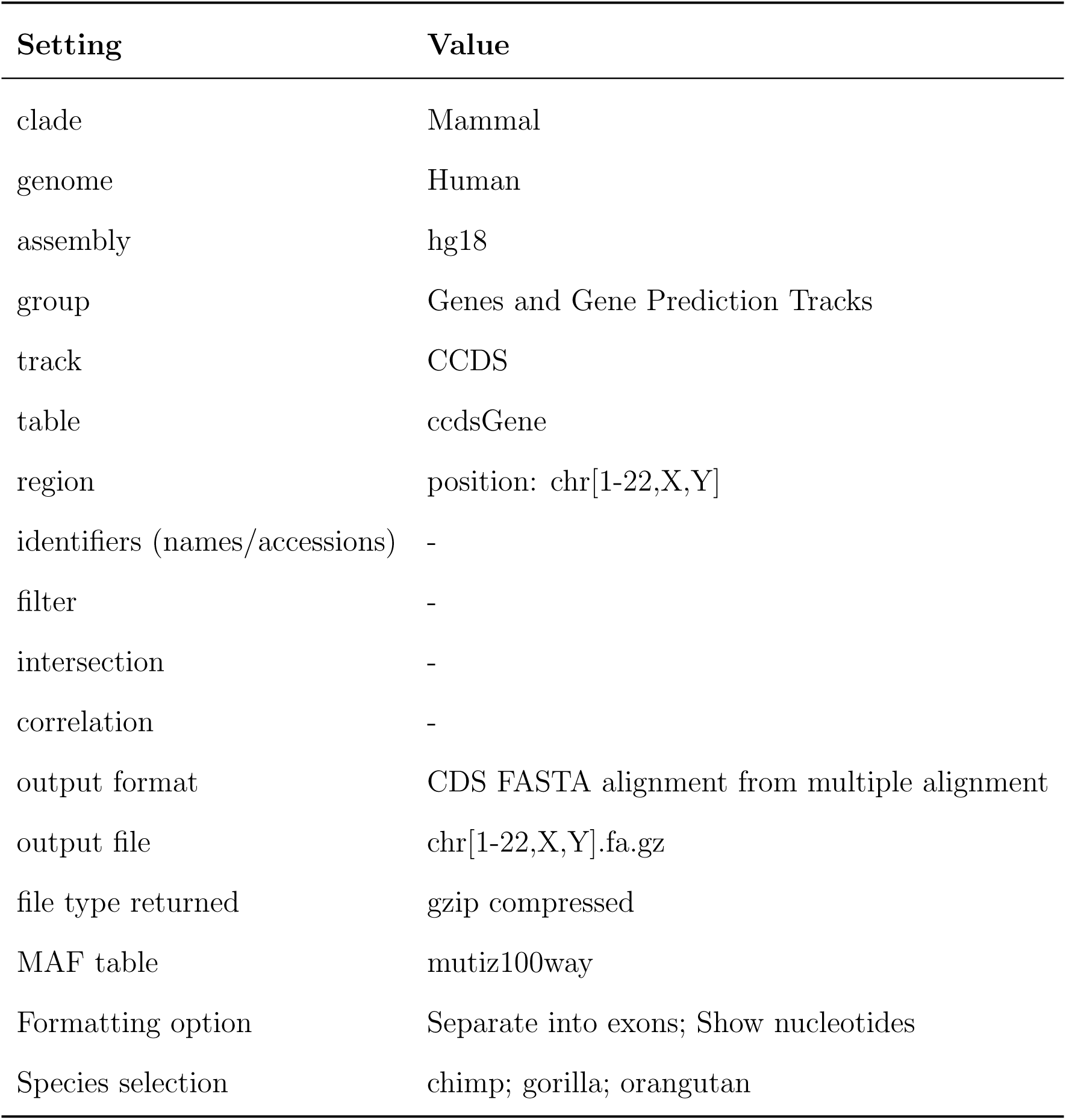

The CCDS alignments were filtered according to the following requirements:

- all treated species need to be present in the alignment;
- divergence from human reference has to be below 10%;
- start codons need to be conserved;
- stop codons need to be conserved;
- no frame-shifting gaps are allowed;
- no gaps longer than 30 bases are allowed;
- no premature stop codon is allowed;
- exon lengths need to be longer than 21;
- genes need to have the same number of exons in all species.

This was done with:

. / FilterMSA. py hg18*-*unfiltered. msa. gz 4 hg18*-* filtered. msa. gz

## Intraspecies Data — Variation in Populations

Polymorphism data was downloaded from http://biologiaevolutiva.org/greatape/ (Prado-Martinez et al. 2013) in Variant Call Format (VCF):

server=” publicprimate@biologiaevolutiva. org”

wget *–*c ftp://${server}/VCFs/SNPs/Gorilla.vcf.gz

wget *–*c ftp://${server}/VCFs/SNPs/Homo.vcf.gz

wget *–*c ftp://${server}/VCFs/SNPs/Panpaniscus.vcf.gz

wget *–*c ftp://${server}/VCFs/SNPs/Pantroglodytes.vcf.gz

wget *–*c ftp://${server}/VCFs/SNPs/Pongoabelii.vcf.gz

wget *–*c ftp://${server}/VCFs/SNPs/Pongopygmaeus.vcf.gz

The VCF files need to be index with **tabix** so that SNPs can be found in reasonable time, e.g.:

tabix *–*p vcf Homo. vcf. gz

The population data spans 79 wild- and captive-born individuals to high coverage including all six great ape species which are divided into a total number of 12 populations; the number of individuals per population ranges from 1 to 23 (cf. table S1).

**Table S1.** Number of samples per population (data from Prado-Martinez et al. 2013).

| Population | Samples |
| --- | --- |
| *Homo sapiens* — Non-African | 6 |
| *Homo sapiens* — African | 3 |
| *Pan troglodytes ellioti* | 10 |
| *Pan troglodytes schweinfurthii* | 6 |
| *Pan troglodytes troglodytes* | 4 |
| *Pan troglodytes verus* | 4 |
| *Gorilla beringei graueri* | 3 |
| *Gorilla gorilla diehli* | 1 |
| *Gorilla gorilla gorilla* | 23 |
| *Pongo abellii* | 5 |
| *Pongo pygmaeus* | 5 |

The SNPs are aligned to the *hg18* human reference genome. A library has been written that can extract the data from the VCF files so that polymorphism data can be incorporated to the filtered CCDS alignments. This library is called *libPoMo* and is distributed together with PoMo (https://github.com/pomo-dev/PoMo); documentation can be found at (http://pomo.readthedocs.org/en/v1.1.0/). The PoMo v1.1.0 release was used throughout this paper.

## Assembly of Inter- and Intraspecies Data

The following command assembles the inter- and intraspecies data. It only extracts 4-fold degenerate sites from the remaining CCDS alignments.

*# –s : only print synonymous bases*

MSAtoCounts. py *–*s hg18*–*filtered. msa. gz \

Gorilla. vcf. gz Homo_afr. vcf. gz Homo_non_afr. vcf. gz *\*

Pan_paniscus. vcf. gz Pan_troglodytes. vcf. gz Pongo_abelii. vcf. gz *\*

Pongo_pygmaeus. vcf. gz *\*

hg18*–*all. cf. gz

The output file in counts format hg18-all.cf was used for all analyses.

## One Individual per Population

A standard HKY model was used to infer species trees for data from one individual picked out of each population (HyPhy with nearest neighbor interchange to maximize the likelihood). The data preparation is the same as the one from above (cf.). Only the -i switch is additionally used for MSAtoCounts.py so that it extracts only one randomly chosen individual out of each VCF file.

## Parameter Estimation Results

The repeated analysis shows on the one hand that the ten trees inferred by PoMo only bin into 3 very similar topologies (see Fig. 7a in the main text). The only differences in these are the relationships of the *Pan troglodytes* clade. The inferred transition to transversion ratio *κ* shows low standard deviation and the log likelihoods are also very stable (table S2).

On the other hand, the HKY model applied to randomly sampled individuals infers 6 different topologies for 10 runs (see Fig. 7b in the main text). The relationships of both, the *Gorilla* and the *Pan troglodytes* clades remain unclear. The transition to transversion ratio is slightly lower and has higher standard deviation. Also the log likelihoods vary significantly (table S2).

The HKY model applied to the consensus sequence is expected to be more stable. The 10 replicated simulation runs in general conform these expectations (table S2). Three different topologies are inferred but only the *Chimpanzee* clade mixes. The topology of the *Gorilla* clade differs from the one estimated by PoMo and the one that is presented in the original paper.

In general, we find that PoMo infers shorter relative branch lengths of external branches. This is due to the fact, that polymorphisms are allowed to be present in the populations while standard DNA models such as HKY interpret each observed change as a substitution event. Hence, the likelihood of short branches and fast changes is relatively higher and shorter branch lengths are favored.

**Table S2.** Mean and standard deviations of the transition to transversion ratio (*κ*), the tree heights and the log likelihoods (log *L*) for PoMo, HKY on randomly chosen individuals (HKY) and HKY on consensus sequence (HKY-Cons).

| | $\kappa$ | tree height | $\log L$ |
| --- | --- | --- | --- |
| PoMo | $7.42 \pm 0.0076$ | $0.22 \pm 0.001$ | $-4920324 \pm 240$ |
| HKY | $5.97 \pm 0.08$ | $0.011 \pm 3 \cdot 10^{-5}$ | $-4757326 \pm 4435$ |
| HKY-Cons | $6.46 \pm 0.005$ | $0.01 \pm 2.1 \cdot 10^{-5}$ | $-4663810 \pm 1061$ |

